# On the benchmarking of clustering algorithms and hyperparameter influence for cell type detection in single-cell RNA sequencing data

**DOI:** 10.1101/2025.08.20.671270

**Authors:** Aleksandra Szmigiel, Ivan Gesteira Costa Filho, Ricardo Jose Gabrielli Barreto Campello

**Affiliations:** Department of Mathematics and Computer Science, University of Southern Denmark, 5230 Odense, Denmark; Institute for Computational Genomics, Joint Research Center for Computational Biomedicine, RWTH Aachen University Medical School, 52074 Aachen, Germany

**Keywords:** single-cell RNA-seq, benchmarking study, methodology, graph-based clustering, cell type detection, community detection

## Abstract

Clustering single-cell RNA-seq (scRNA-seq) data and related protocols remains a major challenge due to high dimensionality, sparsity, and noise. Despite numerous benchmarking studies aiming to identify the most suitable clustering methods, many suffer from methodological flaws that can undermine their conclusions. A major challenge in benchmarking is selecting representative datasets that cover the diversity of scRNA-seq experiments and include laboratory-verified labels for reliable evaluation. Consistent preprocessing of all inputs to benchmarked algorithms is crucial, as it significantly impacts performance. Beyond selecting an algorithm, a thorough exploration of hyperparameters is also essential to assess robustness and identify configurations that maximize performance. We focus on proposing an improved benchmarking framework that addresses common methodological issues in prior studies. We illustrate our proposed methodology in a case study comparing the classic Leiden and Louvain clustering algorithms with extensive hyperparameters exploration on a carefully curated collection of real gold standard datasets. By evaluating clustering performance across different hyper-parameter selection scenarios, we show that benchmarking results can be misleading, either overestimating or underestimating performance depending on how the hyperparameter space is explored. In our illustrative case study, benchmarking results do not reveal any practically relevant performance differences between the Louvain and Leiden algorithms. In contrast, we show that overlooked factors such as graph construction and quality functions critically influence clustering outcomes, particularly under suboptimal settings of numerical hyperparameters—the neighborhood size *k* used for similarity graph construction and the resolution hyperparameter in graph-based clustering algorithms. While noticeable trends have been observed in terms of how different (dis)similarity functions affect performance, the impact of this choice is limited and, to some extent, overridden by the graph-building approach. Across different graphs, there is a noticeable trade-off between achieving optimal performance with ideally tuned numerical hyperparameters and maintaining robustness under more realistic, unsupervised, and suboptimal settings. All in all, the analysis of our illustrative benchmarking case study offers clear guidance and objective recommendations for practitioners in the field. Most importantly, as the main contribution of this manuscript, our proposed framework sets a foundation for more reliable scRNA-seq clustering evaluation and benchmarking in future studies.

## 1 Introduction

Single-cell RNA sequencing (scRNA-seq) has become crucial for unraveling the complexity of biological systems on an individual cell level. With the development of sequencing technologies, it is now possible to analyze millions of cells in a single experiment [1].

The goal of clustering of scRNA-seq datasets is to identify cellular populations with similar expression profiles that help explain the heterogeneity within the data [2, 3, 4, 5]. As an unsupervised (or semi-supervised) method, clustering is inherently challenging. In scRNA-seq applications, this complexity is further amplified by the unique characteristics of the data. For instance, scRNA-seq datasets may now include up to 60,000 genes and over a million cells [6, 7], creating challenges related to high dimensionality and computational scalability. Further complexity arises from hierarchical cell relationships [8, 9] and the inherent noise and sparsity of the data [3].

Community-detection clustering [10] has become increasingly preferred for applications on sequencing data, mainly due to the robustness of sparse similarity graph representations as an alternative to high-dimensional feature-vector data descriptors. Currently, widely used community-detection-based clustering packages for scRNA-seq data and related protocols commonly follow a similar pipeline, which consists of the following standard steps as applied on properly preprocessed feature-vector representation of the raw data [11, 12, 13, 14, 15]: *k* -nearest-neighbors-based graph construction, optional edge weighting, and clustering with algorithms such as Louvain [16] and Leiden [17].

Most benchmarking studies on scRNA-seq data clustering [18, 19, 20, 21, 22, 23, 24] focus on comparing these types of *clustering packages*/*tools*, which typically implement entire workflows. In this work, we use “clustering tool” and “clustering package” interchangeably, referring to pipelines that combine upstream processing with downstream analysis, such as clustering, visualization, or differential expression testing. The very initial data preprocessing stage typically includes dimensionality reduction (such as gene selection and Principal Component Analysis—PCA [12, 14]) prior to any graph representation of the resulting preprocessed feature-vector data. These benchmarking studies comparatively assess each package’s performance using a mix of real and simulated datasets across a range of evaluation measures.

Benchmarking studies on scRNA-seq clustering may exhibit critical issues that hinder reliable comparisons between clustering algorithms or their hyperparameter configurations. One important issue follows from the aforementioned focus on comparing packages consisting of end-to-end pipelines, whose integrated treatment obscures the impact of individual steps. For instance, the graph-building and associated edge-weighting steps are often underexplored, despite their potential to significantly affect results. From the lens of the widespread monolithic view of clustering packages in this field, the influence of these steps cannot be trivially dissociated from the effects of the clustering algorithm itself or the configuration of its hyperparameters. In addition, by comparing clustering packages (rather than algorithms or their hyperparameters), where each package applies a different strategy for preprocessing of raw data and/or a different graph representation of the data prior to clustering, such comparisons can be difficult to interpret. Indeed, it can be argued that different data preprocessing strategies not only could substantially influence the results, they effectively mean clustering performance is actually being compared on different datasets.

Hyperparameter selection is another common issue. Relying on a package’s default values, “blind” guessing, or tuning hyperparameters to match the number of clusters in referential datasets with ground-truth labels may undermine opportunities for a more comprehensive comparative evaluation in which aspects such as hyperparameter sensitivity or hyperparameter optimal settings are also assessed and whereby multifaceted insights such as robustness, average (expected) performance, or best performance (full potential) can be unveiled.

An additional key factor for any benchmarking study is the choice of datasets. Ideally, a sufficiently diverse dataset collection should be adopted representing different organisms, protocols, and scenarios. If available, laboratory-produced ground-truth labels can be independently used to assess whether clustering methods recover previously known, biologically meaningful groups. In contrast, the common practice of relying on a very limited number of datasets, and/or using labels somehow generated with the aid of clustering algorithms as reference annotations, may jeopardize the interpretability and reliability of the evaluation in the corresponding studies.

Our *main focus* is on proposing an improved benchmarking methodology designed to address the limitations of prior studies. In order to *illustrate* the effective application of the proposed methodology in practice, as a *secondary, additional contribution* of this paper we also describe a case study involving the benchmarking of two widely used community detection algorithms, namely, Louvain and Leiden, which are both implemented in numerous software packages [14, 12, 25, 15, 11, 13]. We deploy our benchmarking pipeline to explore the hyperparameter space of these algorithms as well as of critical steps preceding their application, noticeably, (dis)similarity computation and graph construction (Figure 1a). We evaluate the results under three distinct scenarios, each defining a different treatment of hyperparameters (Figure 1b). These scenarios, which are reflective of different settings researchers typically individually adopt to perform benchmarks in single-cell RNA sequencing clustering, will help us illustrate that relying on a single setting does not provide a complete picture of algorithm performance or hyperparameter behavior.

**Figure 1:**
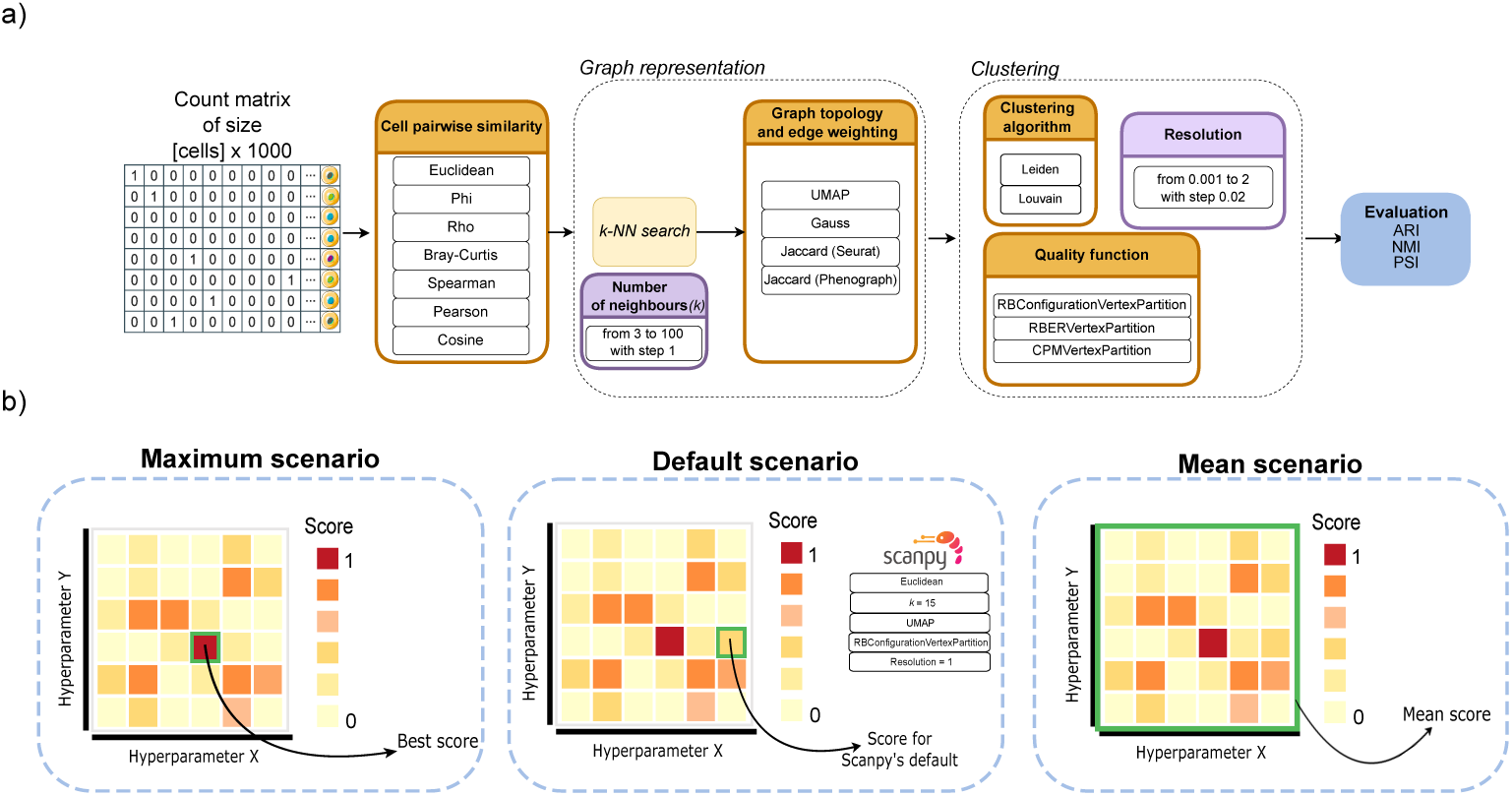
Schematic summary of the proposed benchmarking pipeline as materialized in our illustrative case study: a) We vary non-numerical hyperparameters (Similarity measure, Graph topology and weighting, Quality function for Leiden and Louvain) as well as numerical hyperparameters (Graph neighborhood size *k*, Resolution for Leiden and Louvain); b) The evaluation is performed under three scenarios. The Maximum scenario represents the best score achieved across all tested hyperparameter combinations, reflecting the performance ceiling. The Default scenario uses the default Scanpy settings and captures typical user performance. The Mean scenario reports the average performance across all hyperparameter combinations, simulating uninformed or random selection.

We find that the performance differences between the algorithms are not practically relevant across a collection of real, gold standard scRNA-seq datasets. This is an interesting result in itself, as it does not support the prevailing belief that Leiden is superior, given that it builds upon Louvain. However, the graph construction step prior to applying the algorithms plays a critical role in the clustering outcome. For instance, in Scanpy [12], we observe that the Gauss edge weighting scheme, in which edge weights are computed using a Gaussian kernel [26][27], yields the best results under optimal hyper-parameter selection. In contrast, the UMAP-based graph weighting [28] tends to produce more robust and stable results overall. Additionally, we find that different quality functions optimized by Louvain and Leiden perform best in distinct resolution ranges and differ significantly in their robustness to the resolution hyperparameter.

## 2 Literature review: methodological gaps in scRNA-seq data clustering benchmarking studies

Benchmarking studies on scRNA-seq data clustering use diverse methodologies, ranging from straightforward comparisons of ready-to-use tools to evaluations organized by algorithm type or dataset characteristics. These studies frequently conclude that no universal best tool exists, underscoring the need for evaluations considering dataset-specific performance, whereby graph-based clustering methods are generally reported to yield more satisfactory results when applied to sparse and high-dimensional datasets [18, 19, 24, 20, 21, 29, 22]. The results of existing benchmarking studies should be interpreted with caution due to common methodological gaps, most noticeably, inconsistent preprocessing and limited dataset or hyperparameter selection, as summarized in Table 1 and discussed in the following sections.

**Table 1:**
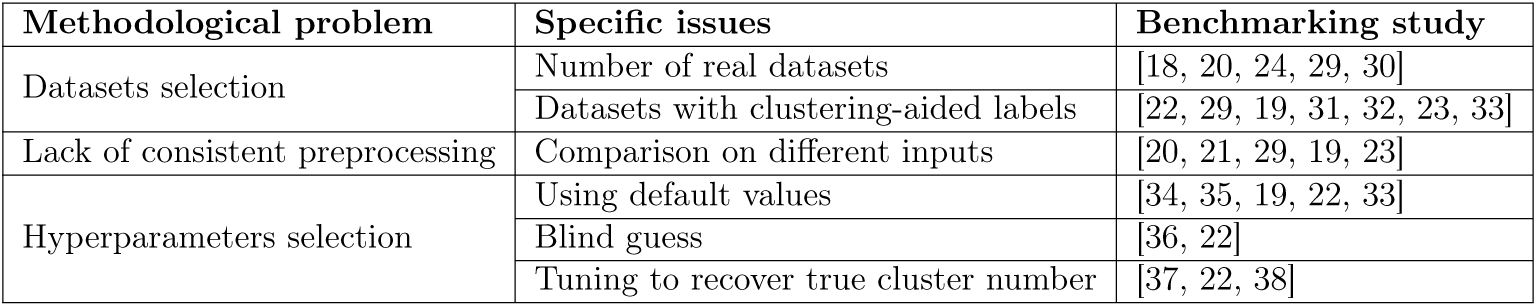
Summary of methodological problems in benchmarking studies.

### 2.1 Datasets selection

One issue in benchmarking scRNA-seq clustering methods is the choice of datasets used for evaluation. Reliable assessment of clustering performance requires datasets with known ground-truth labels. In scRNA-seq, datasets are generally classified as gold or silver standard [39, 40, 35]. Gold standard datasets contain cell types identified through orthogonal methods, rather than gene expression alone. Techniques such as fluorescence-activated cell sorting (FACS) or image-based methods like smFISH provide independent validation of cell types [41]. Reliable gold standard labels can also come from well-characterized sources like distinct cell lines, PBMCs, or early embryonic cells. However, these datasets often lack the complexity of heterogeneous tissues and are typically too small for evaluating large-scale methods [3].

Due to the challenges of generating gold standard labels, researchers often also rely on silver-standard datasets, where cell types are assigned using computational approaches. Most often, this involves clustering followed by annotation with marker genes [42]. There is a multitude of studies that evaluate the performance of clustering algorithms using dataset labels that were originally generated with the aid of clustering algorithms [22, 29, 19, 31, 32, 23, 33] (see Supplement—Evaluation with clustering-aided labels). The potential issue is that, if a clustering algorithm helped produce the labels, then similar algorithms may perform better on those datasets only by matching the particular characteristic captured during the label creation process.

Another issue is the limited number of real datasets often used for bench-marking, since reliable evaluation requires diverse datasets that span various biological contexts. For instance, Duò et al. [24] used only four real datasets in their study, whereas Lin et al. [30] evaluated the performance of their own method on three real datasets. Some studies create multiple datasets by sampling subsets from one or two large datasets under different settings, such as varying the number of cell types or cells [18, 20]. While this introduces variation, it does not replace true biological diversity.

A common strategy to compensate for a limited number of real datasets in benchmarking studies is the reliance on synthetic datasets [30, 24]. However, most simulators struggle to capture complex experimental designs without introducing artifacts, which can result in overly optimistic performance estimates [43]. Although synthetic data allows for a desirable control over certain properties, conclusions based on such datasets should be treated with caution and considered complementary to results from real data.

### 2.2 Inconsistent preprocessing

Clustering tools for scRNA-seq data typically involve preprocessing steps such as normalization, gene or cell filtering, and dimensionality reduction. These steps usually vary widely across software packages. Benchmarking studies that include tools with different preprocessing pipelines [23, 21, 29, 19, 20] make it difficult to draw reliable conclusions about clustering performance, as the compared results are ultimately produced from datasets that can be fundamentally different (despite having been produced from a common raw data source). Differences in normalization can alter cluster structure (e.g., by changing co-variances), while distinct dimensionality reduction methods will yield inputs with varying information.

For example, Krzak et al. [23] evaluated 13 clustering tools, noting that eight incorporated additional preprocessing, but only four of these allowed users to disable it. Nine tools let users choose the number of dimensions for analysis, while just five allowed selection among dimensionality reduction methods like PCA, t-SNE, or UMAP, and not all five supported all options. These differences in functionality can make direct comparisons between clustering algorithms more challenging.

A meaningful comparison requires that all applicable algorithms be tested across the same preprocessing and hyperparameter settings. Nasrollahi et al. [29], for instance, compared Seurat’s [14] default preprocessing pipeline with alternative preprocessing configurations. However, while the Leiden and Infomap clustering algorithms were tested across the full range of preprocessing variants, the Louvain algorithm was evaluated only on the default configuration, despite being also compatible with the other variants. Such differences in experimental design may affect the comparability of results. When algorithms support the same preprocessing options, evaluating them under the same shared conditions will lead to more consistent comparisons.

### 2.3 Hyperparameters selection and implementation differences

Many benchmarking studies select hyperparameters using either default settings or blind guesses [19, 34, 36, 22]. This approach can be problematic, particularly in light of the inconsistencies in preprocessing discussed in Section 2.2. Default settings are often optimized for idealized conditions that may not reflect real-world data, while blind guesses are difficult to justify or generalize across datasets in an unsupervised learning setting.

Another common approach is adjusting hyperparameters to recover a known number of clusters. For instance, Watson et al. [37] adjusted the resolution parameter until the number of clusters matched the annotations or until 1,000 iterations were reached. However, since the main goal of scRNA-seq clustering is to discover unknown cell types or subtypes, this strategy does not help in choosing hyperparameters when true labels are unavailable. A similar issue appears in the study by Liang et al. [22], where resolution was adjusted to match the true cluster number for each dataset. Additionally, the ground-truth labels for some datasets were generated using Seurat, and Seurat was one of the benchmarked clustering tools in that study. In such cases, tuning Seurat’s resolution to reproduce its own outputs may introduce bias and make interpretation of comparative results challenging.

Hyperparameter values vary across popular clustering packages. For example, Seurat sets the default number of neighbors for *k* NN graph construction to 20, while Scanpy uses 15. Even with the same *k* value, the actual number of neighbors can differ due to implementation details. For instance, Seurat includes the query point in the search and counts it towards *k*, while Scanpy includes it during the search but removes self-loops, yielding *k −* 1 neighbors. PhenoGraph [25] excludes the query point entirely. These differences lead to distinct graph topologies (Figure 2a).

**Figure 2:**
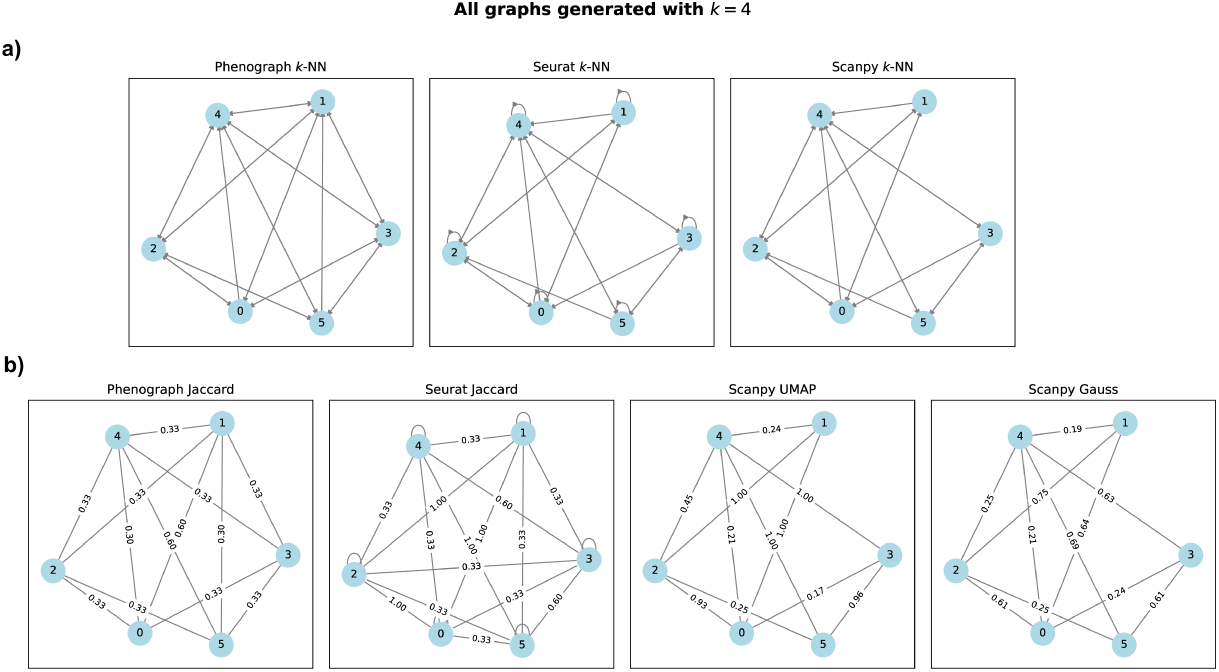
Differences in a) *k* NN graph topologies and b) edge weighting across Scanpy, Seurat, and PhenoGraph. a) There are differences in the *k* NN graph construction step between the packages. b) After application of edge weighting schemes to *k* NN graph, there are even more variations in both the topology (due to removal of zero weight edges and/or creation of new edges to accommodate non-zero weights between non-adjacent nodes) as well as in the edge weights.

Furthermore, Seurat computes approximate nearest neighbors by default, whereas in Scanpy this behavior depends on dataset size, with exact *k*NN computed for smaller datasets.

Another challenge in hyperparameter selection is the lack of standardized terminology across clustering workflows. For example, Seurat’s documentation describes its output as a Shared Nearest Neighbor (SNN) graph built from a *k* - nearest neighbor (*k*NN) graph, where edge weights are computed using Jaccard similarity between the corresponding end points (vertices/nodes). However, the resulting graph topology differs from the original SNN definition by Jarvis and Patrick [44], which requires mutual inclusion in each other’s *k* NN lists for two points to be considered shared neighbors. In contrast, PhenoGraph [25] also applies Jaccard similarity but does not refer to its output as an SNN graph. This method calculates Jaccard similarity between two points only if one is in the other’s neighborhood, resulting in an initially asymmetric graph that is later symmetrized (Figure 2b). Although this approach may more closely resemble the original definition of SNN, it still does not fully meet the criteria of mutual nearest neighbors (see Supplement—Jaccard similarity in Seurat and Phenograph).

## 3 Materials and methods

### 3.1 Data collection

The datasets used in our illustrative benchmarking study were derived from different organisms, protocols, and laboratories. We aimed to find datasets that provided non-normalized cell counts, but in some cases, those were not available. As we advocated above in this paper, our collection consists primarily of gold standard datasets in which cell type labels were determined through laboratory methods or based on well-defined developmental stages. It is worth noticing that our collection also contains a subgroup not fully aligned with a strict definition of gold standard in which datasets are ideally annotated using orthogonal methods not reliant on sequencing [3]. In this subgroup, although clustering was not used to assign labels, alternative computational approaches may have been employed.

Our collection of datasets includes as many datasets as possible used in various benchmarking studies outlined in Section 2, where it could be confirmed that labels were produced without any form of clustering aid. A summary of the datasets is shown in Table 2, while a detailed description is provided in Supplement—Datasets description.

**Table 2:**
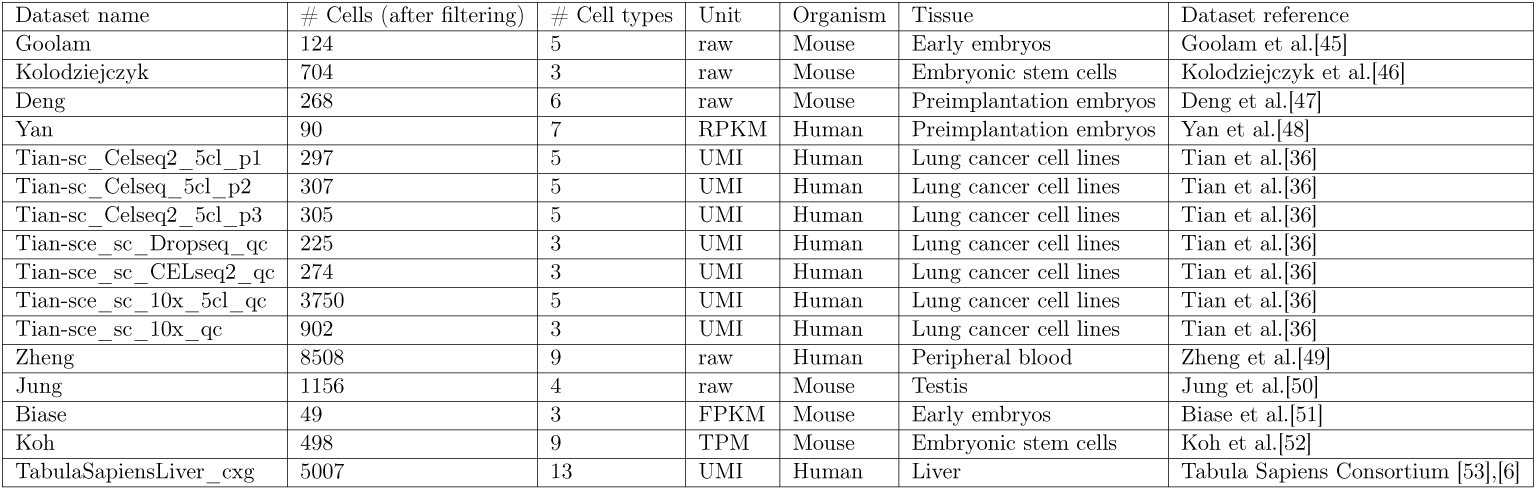
Overview of single-cell RNA-seq datasets used in this study (all datasets were processed to 1,000 genes for downstream analysis).

### 3.2 Data preprocessing

In our analysis, we ensured a consistent preprocessing for all datasets. Since the datasets had been previously processed in various ways by their original authors, we had to adjust our preprocessing pipeline to account for these differences. First, we filtered both cells and genes, adopting the strategy proposed by Watson et al. [37]. We then normalized the counts so that each cell’s total count matched the median total count across all cells, followed by a shifted logarithm transformation. To accelerate analysis, we selected 1,000 highly variable genes for each dataset, following the approach used in the scRNA-seq benchmarking study by Tian et al. [36].

Full details of the preprocessing steps are provided in the Supple-ment—Data preprocessing. It is important to note that there is no single preprocessing workflow that suits all scRNA-seq applications. However, the most important principle for meaningful benchmarking of clustering algorithms is to ensure that *for all models being compared (clustering algorithms and/or combinations of their hyperparameters) each input (dataset) must be exactly the same.* This is achieved by the preprocessing workflow adopted in this paper.

### 3.3 Clustering algorithms

In our illustrative study of the benchmarking methodology proposed in this paper, we focus on the comparative evaluation of two community detection algorithms, Leiden and Louvain, which are among the most widely used in the field and are available in packages such as Scanpy, Seurat, and PhenoGraph.

After the study by Traag et al. [17], the Leiden algorithm has become the favored clustering method in Scanpy, as reflected in its use in official tutorials, while Louvain still remains the default in Seurat. Traag et al. showed that Leiden outperforms Louvain in addressing the issue of disconnected communities, a known limitation of the Louvain algorithm, based upon results from six real-world networks; however, none of these networks were of biological origin, and specifically, none represented gene expression data.

It is important to note that the implementation of the Louvain algorithm differs between Seurat and Scanpy, as described in the work by Rich et al. [54]. In contrast, the Leiden implementation is consistent across both packages. In this study, we utilize the Scanpy implementation of the Louvain algorithm, which is based on the *vtraag-louvain* library [55].

Nashrollahi et al. [29] compared Louvain (using the Seurat implementation) and Leiden in a manner that was criticized above in Section 2.2. The authors stated that Leiden and Louvain will usually produce similar outcomes; however, the benchmarking of the Louvain algorithm was limited to a smaller number of combinations compared to the Leiden algorithm.

Up to this date, and to the best of our knowledge, no study has systematically compared the Leiden and Louvain algorithms across a commonly preprocessed collection of scRNA-seq datasets while exploring the large hyperparameter space that both algorithms—as well as the graph representation required by these algorithms—accept. In this work, we apply the Leiden and Louvain algorithms across a range of hyperparameter configurations.

### 3.4 Hyperparameters selection

As illustrated in Figure 1a, there is a suite of hyperparameters involved across the different steps of our benchmarking pipeline. In the first step, the proximity between cells is computed using a (dis)similarity measure of choice, as a hyperparameter. Considering the results of previous benchmarking studies [56, 37], we evaluate correlation-based measures (`spearman`, `pearson`), Co-sine similarity (`cosine`), three measures based on proportionality (`phi`, `rho`, `braycurtis`), and Euclidean distance (`euclidean`). Euclidean distance was included in the analysis because, despite previous findings showing its inferior performance with scRNA-seq data [37], it remains the default dissimilarity measure in widely used tools such as Scanpy and Seurat.

In high-dimensional data such as scRNA-seq, proximity is more efficiently and effectively represented in the form of sparse similarity graphs, where cells are encoded as nodes and the closest similarities in gene expression are represented as edges. In scRNA-seq applications, *k* -nearest neighbors (*k*NN) variants are the *de facto* choice of similarity graphs. This category includes both unweighted and weighted variants, i.e., graphs that may or may not contain weights on edges, as well as structural variants, e.g., graph topologies based on simple, mutual, or shared nearest neighborhoods. In this study, we considered four different approaches to constructing these graphs. First, we used Scanpy’s own methods—specifically, simple-neighborhood *k* NN graphs with UMAP and Gaussian weighting (hereafter referred to as `umap` and `gauss`, respectively). Additionally, Scanpy provides a wrapper for PhenoGraph’s method for constructing shared-neighborhood *k* NN graphs, which we also considered in our study (`jaccard_phenograph`). Finally, we also evaluated Seurat’s method for shared-neighborhood *k* NN graph construction (`jaccard_seurat`). PhenoGraph and Seurat apply different variants of Jaccard weighting (see Supplement—Jaccard similarity in Seurat and Pheno-graph). As a built-in, default behavior, `jaccard_seurat` relies on approxi-mate *k*NNs, and we retain this design choice in our analysis. Exact *k*NNs are used for `umap`, `gauss`, and `jaccard_phenograph`.

A range of *k* values is considered for constructing the graphs. The starting value is *k* = 3 and it is incrementally increased up to *k* = 100 with a step size of 1. However, for datasets with less samples than the upper limit of *k* = 100, the maximum *k* value is set to 90% of the dataset size (number of cells), rounded up to the nearest integer.

Since not enough attention has been given in the literature to studying how the choice of quality function for the Leiden or Louvain algorithms affects community detection, in this study, we also evaluated three quality functions that incorporate a *resolution* hyperparameter and are available across both the Scanpy and Seurat packages. The equations corresponding to each quality function are provided in Supplement—Quality functions. For each quality function, we compared the resolution hyperparameter across a range from 0.001 to 2, with increments of 0.02, resulting in 100 different resolution values.

### 3.5 Clustering evaluation

For the evaluation of clustering, we relied exclusively on external validation criteria. Specifically, we utilized the well-known Adjusted Rand Index (ARI) and Normalized Mutual Information (NMI). Additionally, we considered the Pair Sets Index (PSI), which is an external validation measure based on pair-set matching [57]. PSI was proposed as an alternative to ARI for assessing clustering results in scRNA-seq data, which penalizes the incorrect clustering of both rare and abundant cell types equally [37]. Following the principle that no single measure can fully capture all possible aspects and perspectives of unsupervised clustering, we adopted the conservative approach of performing experiments and analyzing the three different measures above.

In terms of reporting, we focus mainly on major patterns and observations that are largely consistent across multiple measures. Bearing this in mind and for the sake of conciseness, we primarily report results using ARI in most of Section 4. ARI has been chosen for the following reasons: (i) it is the most widely used measure; (ii) it is less sensitive than PSI to deviations in the number of detected clusters with respect to the ground-truth, which may be over-penalized by PSI irrespective of the overall quality of the compared partitions (see Supplement—Clustering evaluation); (iii) results for NMI were largely consistent with ARI and, therefore, they have been omitted.

## 4 Results

We evaluate the performance of our algorithms across the collection of 16 datasets described in Section 3.1. For the largest dataset in our collection, the Zheng dataset, we were unable to compute the `jaccard_seurat` graph topology because Seurat created graphs with an excessive number of edges, exceeding available computational resources. To navigate the high-dimensional hyperparameter space, we analyze our data through three aggregation strategies. These strategies define how we handle the variable hyperparameters to reach a single score of clustering validation for each fixed target under analysis. A target can be the algorithm itself or a fixed combination of specific categorical hyperparameters:

- Maximum scenario: For each fixed target, we select the absolute maximum score found among the millions of combinations of variable hyper-parameters, per dataset. Tuning all variable hyperparameters simultaneously according to ground-truth labels reveals the ultimate performance ceiling for a given target and dataset, which corresponds to an optimistic result that is not straightforward to obtain in practice given that clustering is an unsupervised task.
- Mean scenario: For each fixed target, we average the scores over the millions of combinations of variable hyperparameters, per dataset. This represents the expected performance under an uninformed, completely random hyperparameter selection.
- Default scenario: This scenario reflects the standard user experience and, technically, it is not an aggregation, but rather an arbitrary selection. For each fixed target, we set all variable hyperparameters to default values in Scanpy (Similarity Measure = `euclidean`, Graph Type = `umap`, *k* = 15, Quality Function = `RBConfigurationVertexPartition`, Resolution = 1). This scenario primarily evaluates the baseline performance for typical users adopting default settings.

### 4.1 Leiden and Louvain perform similarly overall

We first assessed whether there were differences in performance between the two algorithms. We compared their performance distributions for the different clustering evaluation measures across the three previously defined scenarios.

Using the Wilcoxon signed-rank test, we identified statistical differences only in the mean performance scenario, specifically in favor of Louvain. However, as shown in Figure 3, the distributions are nearly identical and overlap significantly. The statistical significance over ranks in that scenario is a result of Louvain’s scores being systematically higher by a negligible margin, typically appearing only at the third decimal place (see Supplementary Table 6). Because the Wilcoxon test operates on relative rankings rather than the magnitude of difference, it captures this consistency without reflecting that the actual change in mean scores is too small to be practically meaningful.

**Figure 3:**
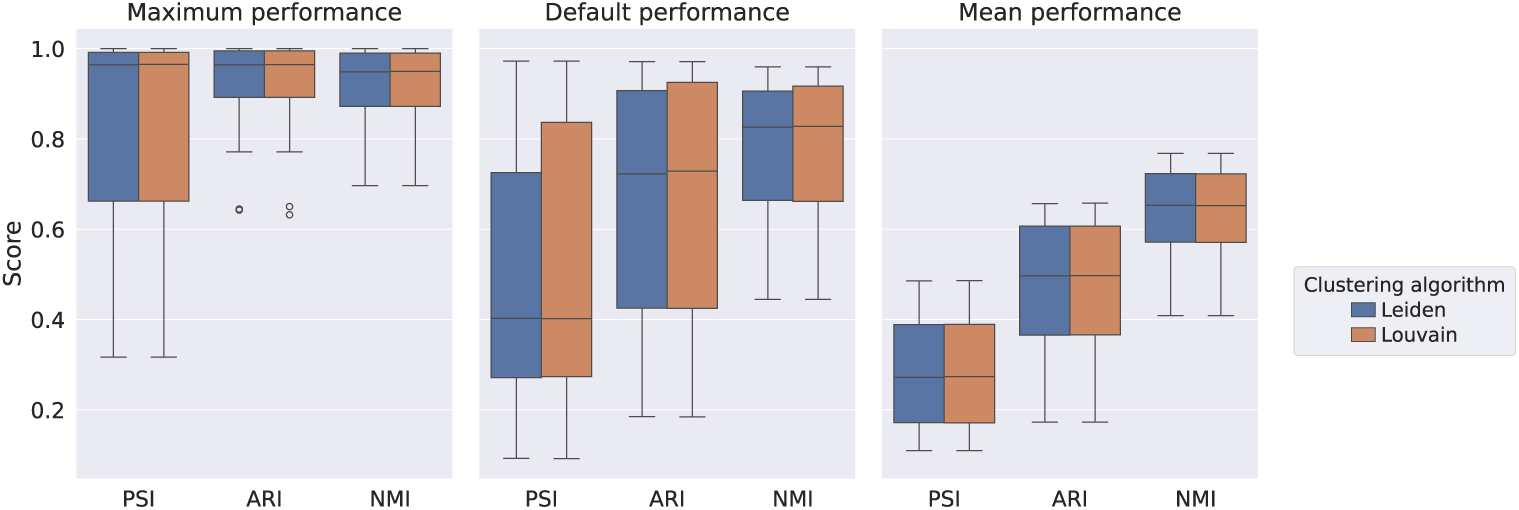
Boxplots of clustering evaluation measures for the Leiden and Louvain algorithms under Maximum, Default, and Mean performance scenarios. Each boxplot summarizes results across 16 datasets.

Visual inspection of the boxplots in Figure 3 shows a difference in the default performance scenario, where the upper quartile of the Louvain boxplot is higher than that of Leiden across all evaluation measures. This effect is primarily driven by a single dataset, in which Louvain outperformed Leiden by approximately 0.2 in ARI and PSI, and around 0.1 in NMI. As this difference is limited to a single case, no general trend of superior performance can be attributed to either algorithm. Bearing this in mind and for the sake of conciseness, in the remainder of our analysis we report results for Leiden only (as the most recent algorithm), noting that the same quality functions are optimized by both algorithms and the main overall conclusions do not change. All corresponding results for the Louvain algorithm are provided in the Supplementary Material.

### 4.2 Clustering performance is dependent on dataset-specific hyperparameter sensitivity

We investigated the behavior of the Leiden algorithm across our collection of datasets to identify patterns in clustering optimization difficulty. Figure 4 highlights the relative “difficulty” of each dataset for the given clustering task.

**Figure 4:**
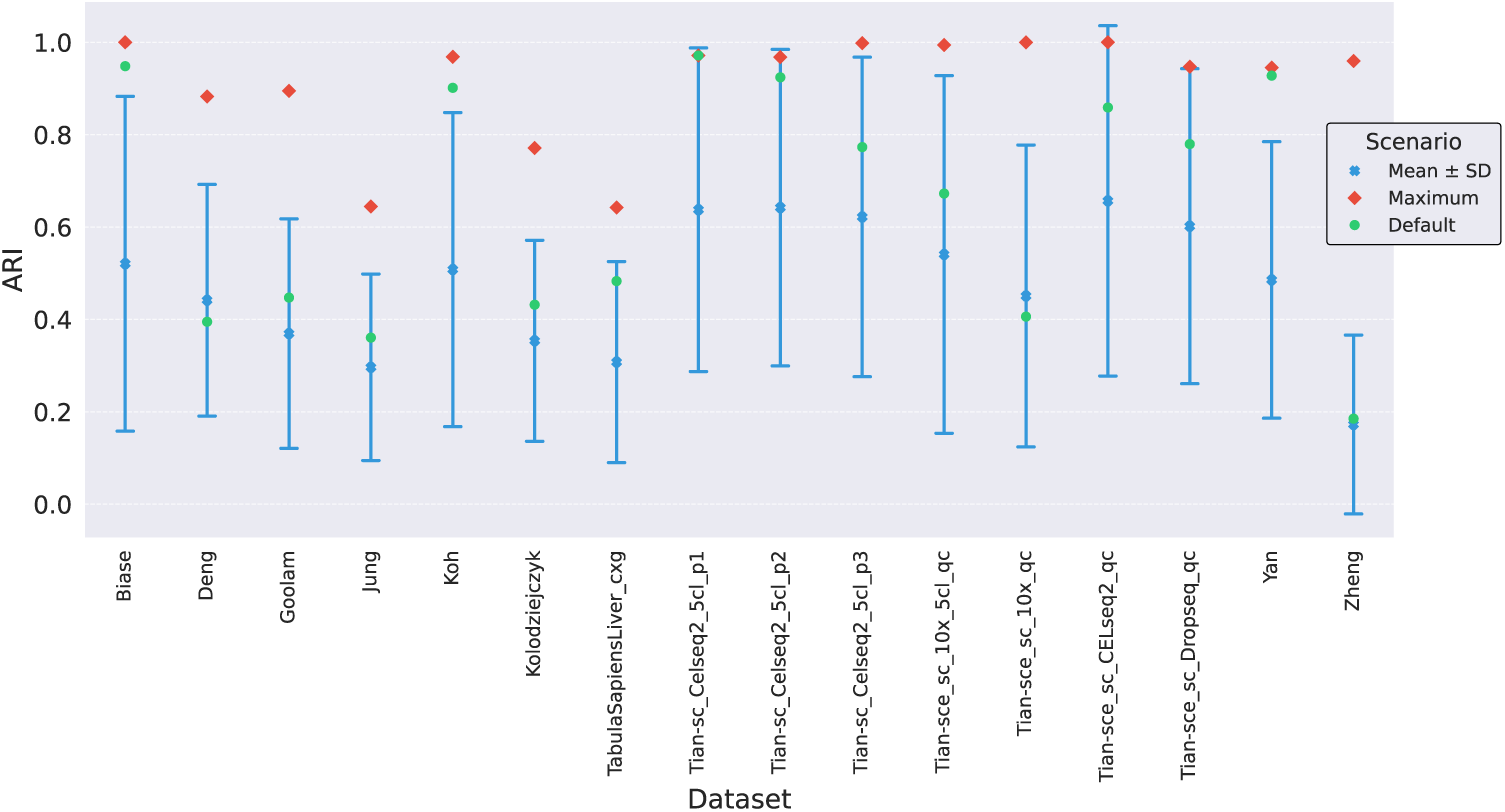
ARI scores per dataset for the Maximum, Default, and Mean performance scenarios. Standard deviation (SD) bars are also displayed for the Mean aggregation scenario.

#### The “Hard” Category

For Zheng, TabulaSapiensLiver_cxg, Kolodziejczyk, Jung, Goolam, and Deng datasets, we observe low mean performance scores, generally below 0.4 (with Deng slightly higher), accompanied by relatively narrow standard deviation (SD) bars. This pattern is evident not only for ARI in Figure 4, but also across all evaluation measures (Supplementary Figures 7, 8). The smaller variability around lower scores indicate that most hyperparameter configurations perform relatively poorly on these datasets. A partial exception is Tian_sce_sc_10x_qc, which exhibits a relatively modest mean performance but with higher variability. Across this group, the maximum scenario performance stands out as an outlier with respect to the expected behavior by chance. The default configuration is not just sub-optimal; it is statistically indistinguishable from a random, uninformed choice. Overall, these datasets require precise, dataset-specific tuning to overcome consistently low performance.

#### The “Intermediate” Category

For Biase, Koh and Yan datasets, we see higher mean performance and wider SD bars as compared to the “hard” category. While the average is still held down by some poor configurations, the Scanpy’s default performs exceptionally well, sitting outside the SD bar. Although the absolute maximum can still improve upon the default scenario, the default already provides a performance level that is not only superior to an uninformed hyperparameter choice, but it is also high in absolute terms. This suggests that the default configuration is well-suited to the structure of these datasets.

#### The “Easy” Category

For the remaining datasets, which consist of the Tian collection with *exception* of Tian_sce_sc_10x_qc, the mean performance is higher and the maximum performance, despite being very high in absolute terms, falls within or near the SD range. On these datasets, a large proportion of the hyperparameter space results in high performance. The optimal region is a broad plateau rather than a localized peak; many hyperparameter configurations lead to a high-quality clustering result.

### 4.3 Clustering is highly dependent on the choice of hyperparameters

To isolate the influence of the non-numerical hyperparameters, we examine the 84 possible combinations of Similarity Measures (7), Graph Types (4), and Quality Functions (3). We handle the numerical hyperparameters (Resolution, *k*) by applying our three established scenarios (maximum, mean, default), effectively collapsing the numerical dimensions into a single representative score for each categorical combination.

To account for the varying levels of dataset difficulty, we calculate the performance of each combination as a mean rank across our collection of datasets, where methods were ranked within each dataset according to their ARI scores. This ranking approach avoids issues that arise when averaging raw scores across “Easy” and “Hard” datasets. For example, a method may achieve high scores on “Easy” datasets and low scores on “Hard” ones; however, the lower scores on “Hard” datasets often reflect a ceiling on achievable performance rather than poor method quality. Directly averaging raw clustering scores can produce a misleading average that masks the method’s true relative performance.

We visualize results as a grid of heatmaps (Figure 5, panels a, c, and e), where each row of the grid represents a scenario, each column of the grid represents a Quality Function, and the individual cells in each heatmap show the mean ranking obtained with the corresponding combination of Similarity Measure and Graph Type. The color scales are normalized independently for each grid row. Consequently, colors represent relative rankings within a single scenario and are not comparable across grid rows.

**Figure 5:**
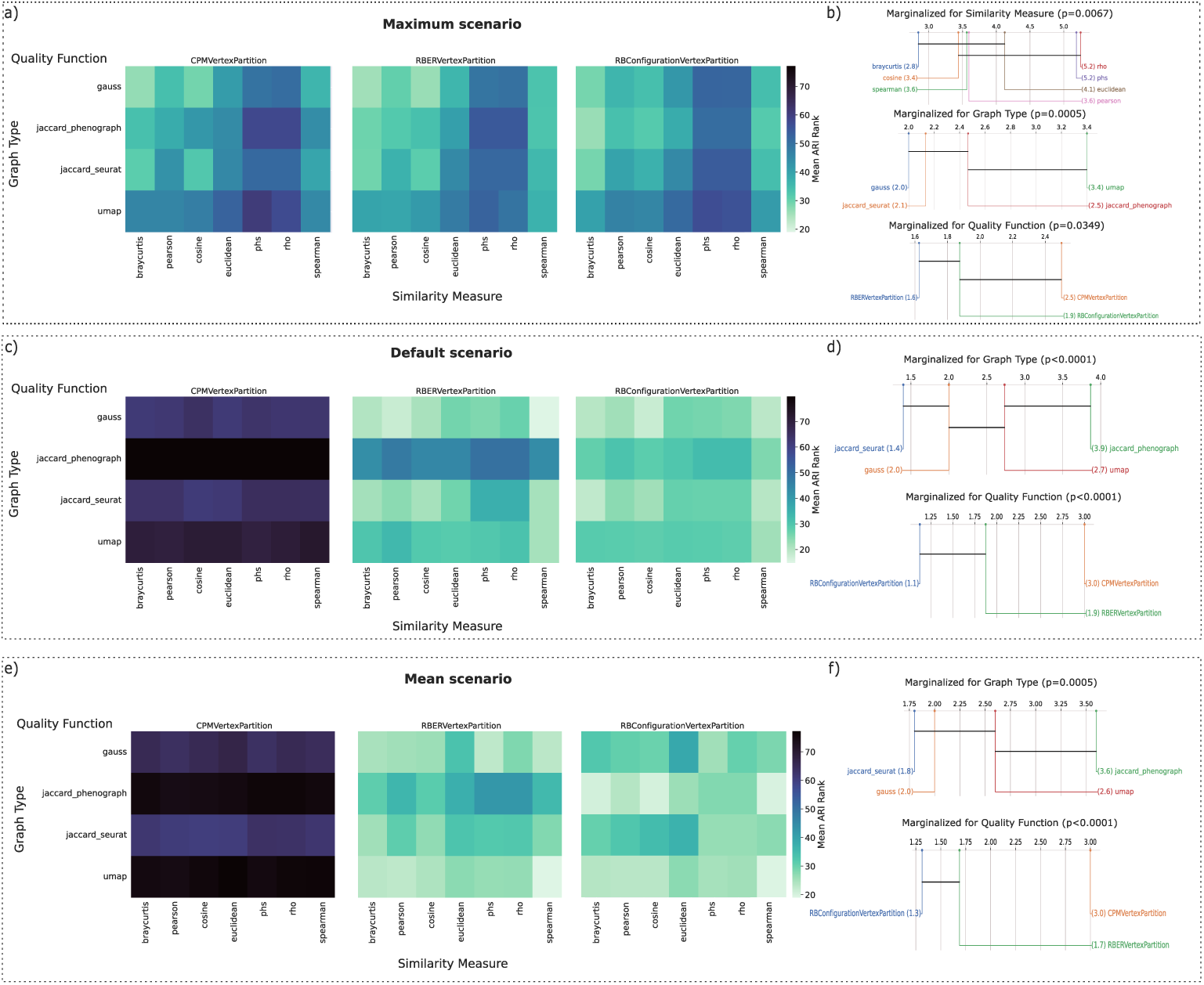
Performance ranking of categorical hyperparameter combinations across three evaluation scenarios. To isolate the effect of non-numerical hyperparameters, we evaluated all combinations of Similarity Measure, Graph Type, and Quality Function, while collapsing Resolution and *k* into three scenarios (Maximum, Default, Mean). Performance is shown as mean rank across datasets. Panels a, c, e display heatmaps for the Maximum, Default, and Mean scenarios, respectively, where each cell represents the mean rank of a Similarity Measure–Graph Type combination for a given Quality Function. Panels b, d, f show critical difference plots for the same scenarios, obtained by marginalizing over other hyperparameters to assess the individual contribution of each category. No critical difference plots are shown for panels d and f for Similarity Measure, as no statistically significant differences were detected in the corresponding analyses.

We use the Friedman test with post-hoc Nemenyi tests to identify statistical significance between all 84 combinations in each scenario (see Supplementary Figure 12 a-c). To complement this global view and isolate the specific contribution of each hyperparameter, we also perform the Friedman-Nemenyi test on each category separately by marginalizing over the influence of the other hyperparameters (Figure 5, panels b, d, and f). In this approach, for each value within a single hyperparameter category (e.g., a specific Similarity Measure), we average the ranks across all other possible settings.

The Friedman test confirms significant performance differences among the 84 categorical combinations across all scenarios, though the Maximum scenario shows no pairwise significance in the Nemenyi test (see Supplementary Figure 12a). This suggests that with optimal numerical tuning (Resolution, *k*), almost any categorical combination can reach a proper score, making Resolution and *k* the most influential factors. In contrast, the Default and Mean scenarios reveal a clear performance gap between combinations in-volving `CPMVertexPartition` and those involving other Quality Functions, indicating that in the absence of numerical hyperparameter optimization, combinations based on this Quality Function underperform relative to the other two (see Supplementary Figures 12b and 12c).

When marginalized across all other hyperparameters, `braycurtis` demonstrates significant superiority over `phs` and `rho` in the Maxi-mum scenario (Figure 5b), a result further confirmed by the heatmaps where `phs` and `rho` consistently occupy the lowest performance tiers regardless of the Graph Type or Quality Function selected (Figure 5a). Notably, the heatmaps for the Maximum scenario re-veal that `braycurtis` paired with `gauss` and `jaccard_seurat`, as well as `cosine` paired with `gauss` and `jaccard_seurat`, consis-tently achieve the highest mean ranks for `CPMVertexPartition` and `RBERVertexPartition`. Supporting the overall strong performance of `braycurtis`, the `RBConfigurationVertexPartition` heatmap shows top mean ranks for `braycurtis` when paired with `jaccard_phenograph`, `jaccard_seurat`, and `gauss`. In contrast, the Default and Mean scenarios show no statistically significant differences when marginalized over Similarity Measure (Figure 5d,f). Consequently, no critical difference plots for Similarity Measures are presented for these scenarios. This indicates that most choices perform comparably under non-optimized conditions, although the mean ranks visualized in the heatmaps suggest that `spearman`, `braycurtis`, and `cosine` tend to achieve slightly higher scores in these scenarios (Figure 5c,e).

Across all scenarios, `gauss` and `jaccard_seurat` consistently rank among the top-performing graph construction methods when marginalized over other hyperparameters, as shown in the critical difference plots (Figure 5b,d,f). In contrast, `umap` performs worst in the Maximum scenario and, together with `jaccard_phenograph`, shows poor performance under the Default scenario for Resolution and *k* (Figure 5c), despite being the recommended default in Scanpy. However, in the Mean scenario (Figure 5e), `umap` paired with `RBERVertexPartition` or `RBConfigurationVertexPartition`, as well as `jaccard_phenograph` paired with the latter, are consistently represented by brighter colors in the heatmap, indicating higher mean ranks compared to other Graph Types in this scenario, and suggesting greater robustness to changes in Resolution and *k*. This robustness is not reflected in the critical difference ranking, as it is offset by the consistently poor performance of both Graph Types when combined with `CPMVertexPartition`.

Finally, marginalization over categorical hyperparameters confirms the strong sensitivity of `CPMVertexPartition` to Resolution and *k*, which is reflected in its underperformance under both the Mean and Default settings (Figure 5d,f). However, it becomes statistically indistinguishable from the second-ranked Quality Function (`RBConfigurationVertexPartition`) under the Maximum setting, when Resolution and *k* are optimized (Figure 5b).

### 4.4 Performance landscapes reveal dataset- and hyperparameter-specific patterns

To bridge the gap between aggregated results in the previous sections and the underlying performance mechanisms, we analyze the interaction between categorical and numerical hyperparameters through the “performance landscapes” (see Figure 6). In these visualizations, we treat the numerical hyperparameters (*k* and Resolution) as the *xy*-coordinate plane, while the ARI score defines the altitude, creating a topography where bright peaks represent regions of optimal clustering whereas dark regions indicate poor performance.

**Figure 6:**
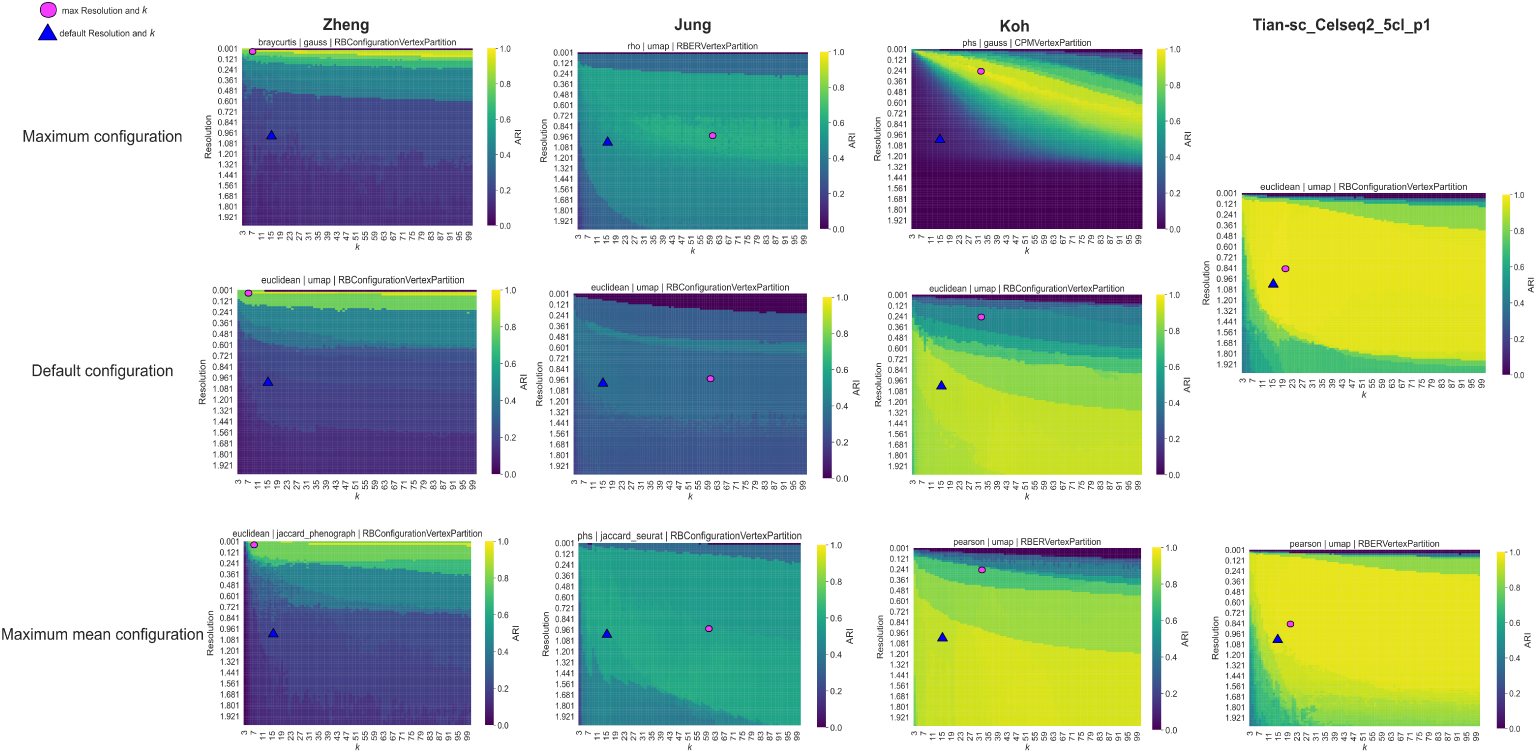
Performance landscapes illustrating the interaction between numerical (*k*, Resolution) and categorical hyperparameters across representative datasets. Each column corresponds to a dataset (Zheng, Jung, Koh, Tian-sc_Celseq2_5cl_p1), and each row represents a different scenario: Maximum configuration (top), Default configuration (middle), and Maximum mean configuration (bottom). Color intensity reflects ARI scores, where bright regions indicate high clustering performance and dark regions indicate poor clustering results. The pink circle marks the optimal numerical coordinates identified in the Maximum configuration, while the blue triangle indicates the default Scanpy settings. Notice that the hyperparameters and results for the 1st and 2nd rows of the 4th column (Tian-sc_Celseq2_5cl_p1) are identical, therefore a single plot has been displayed.

We again evaluate these landscapes across three scenarios: the Maximum configuration (first row of Figure 6), which represents the global peak by selecting the categorical hyperparameters yielding the highest ARI per dataset; the Default configuration (second row of Figure 6), which uses standard Scanpy’s library settings for graph construction and Leiden clustering; and the *Maximum mean* configuration (third row of Figure 6), a scenario that selects categorical settings that maximize the average performance across the entire grid of Resolution and *k* values per dataset.

To further visualize hyperparameters coupling, we fix two static coordinates across all rows: a pink circle representing the optimal numerical coordinates from the Maximum configuration (corresponding to Resolution and *k* giving the overall best ARI), and a blue triangle corresponding to Scanpy’s default values. This reveals the sensitivity of these numerical hyperparameters; a coordinate that is optimal under one categorical regime can fail when the categorical hyperparameters change.

We present results for a selection of representative datasets that serve as examples for the performance categories defined in Section 4.2; the complete set of performance landscapes for all 16 datasets is provided in the Supplementary Material.

The first column of Figure 6 shows the Zheng dataset, a high-difficulty case where the optimal performance region is extremely narrow. This sharp peak explains the large difference between maximum and mean ARI seen in Figure 4. The similarity between the landscapes for every individual categorical settings (see Supplementary Figures 45 and 46) explains the low standard deviation observed around the dataset’s mean performance. Notice that across all three aggregated categorical configurations in Figure 6, the default numerical settings (blue triangle) fall into a sub-optimal range of performance.

Unlike Zheng, which has a hard-to-hit optimum, the Jung dataset (second column of Figure 6) shows inferior results overall, which are limited regardless of hyperparameter choice. While the Maximum mean scenario (third row) succeeds in raising the overall performance floor, no apparent global optimum emerges, suggesting the bottleneck might be the data itself rather than the hyperparameters. This is in conformity with Figure 4, where the mean ARI for this dataset is among the lowest in the study and the gap between maximum and mean performance is relatively smaller.

The Koh dataset (third column of Figure 6) shows a case where the default settings work well (2nd row), with a wide plateau of high performance that covers the default numerical coordinates (blue triangle), suggesting there is some flexibility to vary *k* and Resolution around the default settings without losing performance. This robustness to changes is maintained in the Maximum mean configuration (3rd row); both scenarios use the `umap` graph type, confirming its stability as noted in Figure 5. Notably, the Maximum configuration (1st row), where the absolute maximum (pink circle) is reached, has a much smaller optimal region, indicating a trade-off between peak performance and stability.

Finally, the Tian-sc_Celseq2_5cl_p1 dataset (fourth column of Figure 6) represents the “easy case,” where many different hyperparameter settings achieve an ARI of 1. For this dataset, the default configuration in Scanpy already attains this optimal performance, matching the Maximum scenario. Since these two configurations are identical with the same results for this dataset, only one plot is displayed for both the first and second rows in the fourth column of Figure 6. The Maximum mean configuration also shows a broad plateau of ARI = 1. This indicates that the clustering results remain stable across a wide range of hyperparameter choices for this dataset.

When assessing additional performance landscapes for Graph Type influence, e.g., for datasets such as Biase (Supplementary Figures 15-16), Goolam (Supplementary Figures 19-20), Koh (Supplementary Figures 23-24), Kolodziejczyk (Supplementary Figures 25-26), Tian_sc_Celseq_5cl_p1 to p3 (Supplementary Figures 31 to 36), Yan (Supplementary Figures 43-44), consistent patterns are observed across all Similarity Measures and Quality Functions, with the exception of `CPMVertexPartition`. The `gauss`, `jaccard_phenograph`, and `jaccard_seurat` Graph Types exhibit sim-ilar behavior, characterized by sharper performance changes with respect to Resolution and *k*, and relatively small regions of optimal performance. In contrast, `umap` shows a distinct behavior, maintaining stable performance across broader ranges of both these numerical hyperparameters and a noticeably reduced sensitivity to *k* once Resolution is optimized.

We observe that `CPMVertexPartition` is generally suboptimal, but performs particularly poorly for datasets such as Zheng, TabulaSapiensLiver, Tian_sce_sc_10x_qc and Tian-sce_sc_10x_5cl_qc, where most of the performance landscape is dominated by near-zero ARI values, and only extremely limited regions of higher performance are observed (see Supplementary Figures 45-46, 27-28 and 37-40). Interestingly, all of these datasets were generated using 10x technology, suggesting that experimental characteristics, such as the sequencing protocol, may be associated with specific hyperparameter prefer-ences.

## 5 Discussion and Conclusions

We have observed and characterized a lack of consistency in clustering bench-marking studies as well as the underlying difficulty in drawing reliable conclusions from existing benchmarking pipelines. A common problem is the selection of datasets. Many datasets used in clustering benchmarking studies lack gold standard labels, making clustering results prone to evaluation using referential cell type labels produced with the aid of some form of clustering. In some cases, the same clustering tools that were originally used by the authors of the datasets to assign labels are also evaluated by other researchers as part of a benchmarking investigation. To address this issue, we provide an initial collection of laboratory-verified, gold standard datasets with independently annotated labels, which we have compiled from the literature and systematically preprocessed for benchmarking purposes.

In terms of methodological contributions, to the best of our knowledge, no clustering benchmarking study in single-cell RNA sequencing has simultaneously and systematically considered the influence of multiple categorical hyperparameters, such as the Similarity Measure used to compute cell-cell similarity, Graph Type (which includes graph topology and edge weighting strategies), and the Quality Function optimized by clustering algorithms such as Leiden or Louvain, together with critical numerical hyperparameters such as Resolution and neighborhood size *k*. Our proposed benchmarking framework has been specifically designed to fill this gap.

In terms of results from our illustrative case study intended to showcase the proposed framework in practice, overall, our findings indicate that Louvain and Leiden show highly similar performance, with no substantial differences from a practical standpoint observed across different benchmarking settings. Minor variations are either negligible or dataset-specific and do not indicate a consistent advantage for either method.

We assessed clustering performance from multiple perspectives, highlighting how evaluation results can be misleading unless a thorough and carefully designed experimental analysis is in place. We showed that performance may be underestimated when only a limited portion of the hyperparameters space is explored, while it could also be artificially inflated when focusing on specific datasets that happen to align well with default settings. For instance, the analysis of “Hard” datasets (Section 4.2 and Section 4.4) demonstrates the importance of extensive search: evaluation of only a small number of configurations would likely miss the narrow regions of optimal performance. Such practices risk drawing misleading conclusions about clustering performance.

For a given limited, fixed budget of computational resources, rather than prioritizing comparisons across as many different algorithms as possible, we advocate for a thorough exploration of the hyperparameters space within a selected set of methods. To this end, we evaluate performance under three scenarios that reflect different levels of hyperparameter knowledge (namely, assuming the optimal values were to be known, assuming default values, or assuming a completely uninformed random choice).

We observed a limited impact of the Similarity Measure on clustering performance. While selecting an appropriate measure for a specific dataset may improve results, in general, it is not a dominant factor as compared to other hyperparameters. Overall, `braycurtis`, `spearman`, and `cosine` have shown to be the most reliable choices in our experiments. Although Scanpy’s default `euclidean` distance has been criticized for suboptimal performance in Leiden clustering for scRNA-seq data [37], and is indeed suboptimal to a certain extent according to our general analysis, its overall influence is limited once a similarity graph is constructed. Ultimately, clustering outcomes on these graphs are driven by (topology and) edge weights, which can mitigate differences between Similarity Measures. For that reason, graph construction plays a more substantial role.

According to our findings, different Graph Types exhibit distinct responses to changes in Resolution and *k*, as reflected in their performance landscapes. In particular, `umap` produces broad regions of near-optimal performance, indicating robustness to changes in numerical hyperparameters, whereas other Graph Types tend to have sharper, more localized optima. Maximum performance is often achieved with `gauss`, suggesting that it benefits from careful tuning but is less stable overall. This highlights a trade-off: `gauss` would possibly be preferable could precise optimization be feasible, while `umap` offers a safer, more robust choice under unsupervised uncertainty. Jaccard-based constructions tend to sit somewhere in between.

While there are no obvious winner between `jaccard_seurat` and `jaccard_phenograph` when paired with `RBERVertexPartition` and `RBConfigurationVertexPartition`, clear differences emerge when they are combined with `CPMVertexPartition`: alongside `gauss`, `jaccard_-seurat` is better able to handle the resulting narrow regions of optimal perfor-mance, whereas `jaccard_phenograph` and `umap` show little to no regions of strong performance under this quality function. This behavior is consistent with the overall sensitivity of `CPMVertexPartition` to Resolution and *k*. While it can achieve high performance when these numerical hyperparameters are carefully optimized, its performance under default settings or uninformed random selection is consistently poor. In contrast, `RBERVertexPartition` and `RBConfigurationVertexPartition` show significantly more stable and comparable performance across scenarios, so our findings strongly support these Quality Functions over `CPMVertexPartition`.

Our experiments show that most combinations of categorical hyperparameters can lead to a high-quality clustering result as long as they are associated with strictly optimal values of Resolution and *k*, which could thus be deemed the most influential factors from this standpoint. However, since optimization of these numerical hyperparameters is not possible in practice due to the unsupervised nature of real-world clustering, our results show that the selection of suitable categorical hyperparameters becomes critically important under a more realistic suboptimal choice of Resolution and *k*. Our analyses above are meant to provide objective guidance in this regard.

We noticed that additional characteristics, such as sequencing protocol or tissue of origin, may also influence clustering behavior. In particular, datasets generated using 10x technology were associated with consistently poorer performance for certain configurations in our experiments. Datasets deemed more challenging to optimize (Section 4.2), particularly those derived from embryonic development (e.g., Jung, Goolam, Deng), likely reflect more continuous cellular transitions, which may explain lower clustering scores.

Our work paves the ground for for future extensions along many different directions, for instance, the use of our proposed framework for an independent benchmarking of a broader suite of clustering algorithms, including more recent algorithms claiming to outperform Leiden and Louvain [29]. Another important direction is to investigate how dataset characteristics influence clustering performance, for example through controlled experiments varying sparsity, number of cell types, cell count, or preprocessing choices such as dimensionality reduction. Finally, given the strong impact of graph construction observed in our study, further exploration of alternative graph topologies and weighting schemes remains a promising research pathway.

In summary, we highlight key limitations of current scRNA-seq bench-marking studies and show that insufficient exploration of the hyperparameter space can lead to misleading conclusions about algorithm performance. Building on this, we provide a flexible Snakemake workflow (https://github.com/Campello-Lab/Clustering-benchmarking-for-scRNAseq) that enables systematic and scalable exploration of datasets, algorithms, and hyperparameters, supporting more robust, reproducible, and comprehensive benchmarking.

## 5.1 Key Words

single-cell RNA-seq, benchmarking study, methodology, graph-based clustering, cell type detection, community detection

## 5.2 Authors’ contribution

R.J.G.B.C. and I.G.C.F. supervised the project. Conceptualization and methodology were developed by A.W.SZ., I.G.C.F., and R.J.G.B.C. A.W.SZ curated the datasets for both initial use and later reuse, and implemented the software based on the proposed methodology. A.W.SZ., I.G.C.F, and R.J.G.B.C jointly interpreted the results of the illustrative benchmarking study. A.W.SZ. wrote the original draft, which was reviewed and edited by R.J.G.B.C. and I.G.C.F. Funding was provided by R.J.G.B.C.

## 5.3 Biographical Note

Ricardo Jose Gabrielli Barreto Campello is a Professor in the Department of Mathematics and Computer Science at the University of Southern Denmark. His research focuses on foundational data mining and machine learning, with emphasis on developing new algorithms and mathematical methods for descriptive and predictive analytics.

Ivan Gesteira Costa Filho is a Professor at the Institute of Computational Genomics, University Hospital RWTH Aachen. His work focuses on bioinformatic algorithms and statistical machine learning to study transcriptional, regulatory, and epigenetic mechanisms in cell differentiation and disease.

Aleksandra Szmigiel is a PhD student in the Department of Mathematics and Computer Science at the University of Southern Denmark. Her research involves bioinformatics and the application of machine learning methods to biological data analysis.

## 5.4 Data availability

All benchmarking code, including the improved methodology implemented as a Snakemake workflow, dataset download scripts, and analysis notebooks, is available at: https://github.com/Campello-Lab/Clustering-benchmarking-for-scRNAseq.

