## Supplementary Material for "On the benchmarking of clustering algorithms and hyperparameter influence for cell type detection in single-cell RNA sequencing data"

### 6 Supplement

#### 6.1 Evaluation with clustering-aided labels

In the study by Liang et al. [22], the authors utilized a collection of 10 datasets to assess the performance of 7 clustering algorithms. Three of these datasets were described as "annotated through a typical scRNA-seq analysis workflow", indicating that clustering techniques were applied to identify cell types. Specifically, the datasets with accession numbers GSE180914, GSE171381, and E-MTAB-670 were processed with Seurat by original authors of the datasets, and the resulting cell type annotation was used as a ground-truth for validation of clustering methods in this study. Seurat was also included as one of the benchmarked methods in this analysis and, notably, demonstrated top-tier performance on these datasets across different evaluation metrics employed by the authors.

A similar issue can be found in the study by Nasrollahi et al. [29], where the authors analyzed two real datasets, one of which contained cells annotated using the Seurat pipeline. For the refinement of this annotation, they incorporated the Azimuth tool, which uses reference mapping for annotating cells in scRNA-seq datasets. They retained only the cell types that were consistent across both Seurat and Azimuth annotations. Nonetheless, a potential bias towards the Seurat clustering approach may remain, as it served as the initial method for generating the partition, that the authors treated as the ground-truth. This presents a significant issue, as Seurat is also one of the methods being evaluated in this benchmarking study.

In one of the evaluation studies of scRNA-seq data clustering packages [19], the authors used six real datasets. For instance, the "Mouse brain" dataset (GSE74672) with seven cell types distinguished with divisive biclustering was used for evaluation of clustering tools in this study. Evaluation on this dataset will only allow to conclude about how good the compared clustering packages are in reproducing the behaviour of divisive biclustering. Furthermore, it was shown that Seurat clustering package outperformed other clustering packages on SCP916 dataset. Cell types for this dataset were defined using *FindClusters* function implemented in Seurat. Since Seurat was used to generate the ground truth labels, it is not surprising that it performs well when evaluated against labels prepared with the same method.

Feng et al. [31] used two real datasets for evaluation of dimensionality reduction and clustering tools. One of the datasets [58] that was used for supervised evaluation of clustering, had labels prepared with a clustering method (divisive biclustering). This again highlights a recurring issue: tools that demonstrate high performance on this dataset tend to be those that can best reproduce the behavior of the algorithm originally used to produce the labels.

Studies in which novel tools for scRNA-seq clustering are introduced, are often tested on datasets in which labels were obtained with similar clustering methods. For instance, in the paper by Ren et al. [32], the *Community Detection based on a Stable K-Nearest Neighbor Graph Structure* method proposed by the authors is compared with other clustering techniques. However, the labels for the dataset used in this comparison, sourced from the Human Cell Atlas [59], were obtained via Louvain clustering. Notably, the method proposed by Ren et al. also employs Louvain clustering. The authors compare their method to clustering algorithms of a completely different nature, like density-based clustering. This reliance on labels generated by a specific clustering method introduces bias, undermining the reliability of the comparison and favoring community detection methods that align with the labeling technique.

#### 6.2 Jaccard similarity in Seurat and Phenograph

In Seurat's default workflow, the $k$NN graph is transformed into the output graph by calculating the neighborhood overlap between any two nodes using the Jaccard similarity. Importantly, each node includes itself in its neighborhood when calculating the intersection, which helps distinguish between cases where nodes are direct neighbors (either mutual or one-sided) and cases where they are not neighbors but share common neighbors. This inclusion increases the intersection size when nodes are connected, thus yielding higher similarity scores for actual neighbors. Notably, Seurat's weighting scheme can lead to additional edges in the final topology between nodes that are not direct neighbors but share common neighbors.

Phenograph applies Jaccard similarity in a fundamentally different way from Seurat: the similarity is calculated between point $i$ and point $j$ only if point $i$ includes point $j$ in its neighborhood. Even if they share other $k$-nearest neighbors, the primary condition is that the tested point is present in the other point's neighborhood. This relationship initially is not symmetric—there can be an edge from $i$ to $j$ without a reciprocal edge from $j$ to $i$, if point $j$ does not include point $i$ in its neighborhood. After this calculation, symmetrization is performed, where the edge from $j$ to $i$ is either assigned the same weight as the edge from $i$ to $j$ or half of that weight (in the case of symmetrization by averaging).

Note the difference in edge weight calculation in Figure 2b for the edge between nodes 1 and 2: Seurat assigns a maximum Jaccard similarity of 1, while PhenoGraph assigns a lower weight of 0.33.

**Jaccard Similarity (Seurat):** $N(1) = \{0, 1, 2, 4\}, \quad N(2) = \{0, 1, 2, 4\}$

$$\text{Jaccard}(1, 2) = \frac{N(1) \cap N(2)}{N(1) \cup N(2)} = \frac{4}{4} = 1$$

**Jaccard Similarity (PhenoGraph):** $N(1) = \{0, 2, 3, 4\}, \quad N(2) = \{0, 1, 4, 5\}$

$$\text{Jaccard}(1, 2) = \frac{N(1) \cap N(2)}{N(1) \cup N(2)} = \frac{|\{0, 4\}|}{|\{0, 1, 2, 3, 4, 5\}|} = \frac{2}{6}$$

Another example that highlights the differences between Seurat and PhenoGraph topologies and weighting schemes is the computation of the edge weight between nodes 1 and 5:

**Jaccard Similarity (Seurat):** $N(1) = \{0, 1, 2, 4\}, \quad N(5) = \{2, 3, 4, 5\}$

$$N(1) \cap N(5) = \{2, 4\}, \quad N(1) \cup N(5) = \{0, 1, 2, 3, 4, 5\}$$

$$\text{Jaccard}(1, 5) = \frac{|N(1) \cap N(5)|}{|N(1) \cup N(5)|} = \frac{2}{6}$$

*Note: Even though nodes 1 and 5 are not direct neighbors in Seurat's kNN topology, the similarity is computed based on shared neighbors.*

**Jaccard Similarity (PhenoGraph):** $N(1) = \{0, 2, 3, 4\}, \quad N(5) = \{1, 2, 3, 4\}$

$$N(1) \cap N(5) = \{2, 3, 4\}, \quad N(1) \cup N(5) = \{0, 1, 2, 3, 4\}$$

$$\text{Jaccard}(1, 5) = \frac{|N(1) \cap N(5)|}{|N(1) \cup N(5)|} = \frac{3}{5}$$

$$\text{Final edge weight (after symmetrization)} = \frac{1}{2} \cdot \frac{3}{5} = \frac{3}{10} = 0.3$$

#### 6.3 Datasets description

In the Biase et al. [51] dataset, the cell type labels are likely to be accurate because they were determined at different time points of embryonic development. Another dataset, in which the labels were also determined based on the developmental stages of embryos, is the Deng dataset [47]. Individual cells were dissociated from embryos from different stages of mouse pre-implantation development. Similarly to the previous datasets, the Goolam dataset [45] is another embryonic development dataset that contains cells collected at late

2-cell, late 4-cell, late 8-cell, 16-cell, and 32-cell stages. In the Jung dataset [50], the authors purified cells with FACS protocol. Dataset by Koh et al. [52] consists of FACS purified H7 human embryonic stem cells in different differentiation stages. The dataset of Kolodziejczyk et al. [46] contains mESCs (mouse embryonic stem cells) cultured in three different conditions, making up 3 distinct cell populations. Yan et al. [48] profiled 124 human pre-implantation embryos and embryonic stem cells. This is another example of a dataset with highly confident cell labels because they represent cells from different developmental stages. The Zheng et al. [49] datasets consists of reads collected from FACS-sorted Peripheral Blood Mononuclear Cells.

The next datasets may not fit strictly into the gold standard category because, according to the definition given by Kiselev et al. [3], orthogonal methods for determining cell composition in tissues are those that do not rely on sequencing. In those datasets, clustering wasn't used to produce the labels, however other computational methods were applied. Tian et. al [36] generated datasets, in which ground truth is the cell line identity that can be determined for each cell on the basis of known genetic differences. We also utilize liver cells from the first version of Tabula Sapiens dataset, published in 2022 [53], where cell-level manual annotation was performed by tissue experts using CellxGene browser [6] on a 2D UMAP embedding.

For more details about the collection of datasets, see Table 3.

Table 3: Datasets Information (before processing).

| Dataset | # Cells (before filtering) | # Genes | # Cell types | # Cells per type | Unit | Dataset reference | Source |
|---|---|---|---|---|---|---|---|
| **Gold Standard Datasets** | | | | | | | |
| Goolam | 124 | 41480 | 5 | 6;16;64;32;6 | raw | Goolam et al.[45] | [60] |
| Kolodziejczyk | 704 | 38653 | 3 | 295;159;250 | raw | Kolodziejczyk et al.[46] | [61] |
| Deng | 268 | 22958 | 6 | 49;22;14;37;133<br>13 | raw | Deng et al.[47] | [62] |
| Yan | 90 | 20214 | 7 | 6;12;20;30;16<br>3;3 | RPKM | Yan et al.[48] | [48] |
| sc_Celseq_5cl_p2 | 307 | 14078 | 5 | 107;44;44;71;41 | UMI | Tian et al.[36] | [63] |
| sc_Celseq2_5cl_p1 | 297 | 15564 | 5 | 103;50;50;51;43 | UMI | Tian et al.[36] | [63] |
| sce_sc_Dropseq_qc | 225 | 15127 | 3 | 86;69;70 | UMI | Tian et al.[36] | [63] |
| sce_sc_CELseq2_qc | 274 | 28204 | 3 | 112;81;81 | UMI | Tian et al.[36] | [63] |
| sce_sc_10x_5cl_qc | 3918 | 11786 | 5 | 1191;426;745;828;560 | UMI | Tian et al.[36] | [63] |
| sc_Celseq2_5cl_p3 | 305 | 13426 | 5 | 118;35;41;75;36 | UMI | Tian et al.[36] | [63] |
| sce_sc_10x_qc | 902 | 16468 | 3 | 313;314;275 | UMI | Tian et al.[36] | [63] |
| Zheng | 92043 | 32738 | 9 | 331;6485;191;771;150<br>255;44;53;228 | raw | Zheng et al.[49] | [64] |
| Jung | 1192 | 6354 | 4 | 199;367;298;292 | raw | Jung et al.[50] | GSE113293 [65] |
| Biase | 49 | 25737 | 3 | 20;20;9 | FPKM | Biase et al.[51] | GSE57249 [65] |
| Koh | 498 | 60483 | 9 | 59;76;67;36;51<br>55;22;65;67 | TPM | Koh et al.[52] | GSM2257302 [65] |
| Tabula Sapiens | 5007 | 58482 | 13 | 146;267;421;108;76<br>1558;42;34;1381;245<br>612;83;34 | UMI | Tabula Sapiens Consortium [53] | [6] |

#### 6.4 Data preprocessing

The preprocessing strategy that was applied to datasets was inspired by the study of Watson et al. [37]. First, cells with more than 10% of mitochondrial counts were filtered. Subsequently, the authors of the study filtered cells with less than 200 genes expressed. In our study, two thresholds were applied for filtering genes because the collection of datasets has different numbers of cells and genes. If the number of cells in a particular dataset was in the tens of thousands (Zheng's et al. [49] dataset), cells in which fewer than 1000 genes were expressed were filtered. If the number of cells in a particular dataset was fewer than 10000 cells, cells with less than 100 genes expressed were filtered. In the next step, genes expressed in less than 10% of cells were removed.

Different normalization procedures were applied to datasets, depending on processing applied by the original authors of datasets. In cases where raw counts or Unique Molecular Identifiers (UMIs) were available, they were divided by a size factor. The size factor in single-cell seq RNA normalization can be determined in different ways and should account for sampling variations in a dataset. In our analysis, counts were normalized using the size factor, ensuring that each cell has a total count equal to the median of total counts across all cells. Then a pseudo-count of one was added and a natural logarithm was applied to a dataset, as recommended in the review of best practices for single-cell analysis [5]. For the datasets, that were already normalized (TPM, FPKM), we applied only a shifted logarithm transformation.

To speed up analysis time, 1000 highly variable genes were selected for each dataset, following the approach used in the benchmarking study of scRNA-seq analysis pipelines [36]. For datasets in which raw or UMI read counts were available, a method based on Pearson residuals was applied to extract 1000 highly variable genes. [66]. The analytic Pearson residuals method expects raw, non-normalized input to identify biologically variable genes. For some datasets, raw counts were not available, and in those cases, we applied the dispersion-based method [67].

#### 6.5 Quality functions

In this study, we benchmarked three different quality functions that can be optimized by Leiden or Louvain algorithms.

In Seurat, users can choose a quality function that the developers refer to as *simple modularity.* This function (in Seurat's implementation) includes a resolution hyperparameter that controls the granularity of the resulting clusters. It is important to note that the term *simple modularity* might be mistaken for the original modularity measure introduced by Newman and Girvan [68], which does not involve a resolution hyperparameter. However, in the context of Seurat, *simple modularity* refers to an enhanced version of modularity that incorporates a resolution. This approach was proposed by Jörg Reichardt and Stefan Bornholdt in their 2006 paper [69] (see Equation 1). In this work, Reichardt and Bornholdt modified the original modularity function by introducing a tunable parameter $\gamma$, which adjusts the resolution of community detection. This function is the default in Scanpy, and its name—*RBConfigurationVertexPartition*—is self-explanatory, with *RB* standing for *Reichardt* and *Bornholdt.* Seurat also allows users to apply a so-called *alternative modularity*, which corresponds to the *Constant Potts Model* (Equation 3), also implemented in Scanpy. Understanding these differences in terminology and default settings between packages is important, as they can significantly influence clustering results.

The third quality function that we compare is Reichardt and Bornholdt's Potts model with an Erdős-Rényi null model (*RBERVertexPartition*, Equation 2), which can also be used in Scanpy and includes a resolution hyperparameter.

Equations for all of the quality functions used in this study can be found below:

- Reichardt and Bornholdt's Potts model with a configuration null model (RBConfigurationVertexPartition) [69]:

$$Q = \sum_{ij} (A_{ij} - \gamma \frac{k_i k_j}{2m}) \delta(\sigma_i, \sigma_j) \tag{1}$$

where item $A_{ij}$ is the element of the adjacency matrix (weighted or unweighted), $k_i$ and $k_j$ is the number (weight) of edges to which node $i$ and $j$ are connected, $m$ is the total number of edges (total edge weight), $\gamma$ is resolution parameter, $\sigma_i$ and $\sigma_j$ defined in Kronecker delta $\delta$ denote the community of node $i$ and $j$.

- Reichardt and Bornholdt's Potts model with an Erdős-Rényi null model (RBERVertexPartition) [69]:

$$Q = \sum_{ij} (A_{ij} - \gamma \frac{m}{\binom{n}{2}}) \delta(\sigma_i, \sigma_j) \tag{2}$$

where $n$ is total number of nodes

- Constant Potts Model (CPM) [70]:

$$Q = \sum_{ij} (A_{ij} - \gamma) \delta(\sigma_i, \sigma_j) \tag{3}$$

The term $\frac{m}{\binom{n}{2}}$ in Equation 2 represents *Erdős-Rényi null model*, which is a random graph used as a comparison to the actual graph structure. The concept is inspired by the idea that randomizing the network structure destroys communities, so the comparison between the actual structure and its randomization reveals how non-random the group structure is. In the *Erdős-Rényi null model*, any two vertices have the same probability of being adjacent, ensuring that there is no preferential linking involving special groups of vertices [10].

The term $\frac{k_i k_j}{2m}$ in Equation 1 represents the *configuration null model*, which can be viewed as generalization of *Erdős-Rényi null model* which is a graph designed to match the original one in certain structural features while remaining otherwise random. Edges are rewired randomly, with the constraint that the expected degree of each vertex remains consistent with the vertex's degree in the original graph.

Traag et al. [70] suggested that instead of comparing the network to a random null model, it could be compared to a constant factor, as it has been shown in Equation 3. This quality function tries to maximize the number of internal edges while at the same time keeping relatively small communities [70].

#### 6.6 Clustering evaluation

The abundance of external validity indices for clustering evaluation makes it challenging to select a suitable one. Watson et al. [37] proposed the use of PSI [57], suggesting that it may be less sensitive to the number of clusters and better suited for evaluating clustering errors involving both abundant and rare cell types, in contrast to ARI. However, based on the experiments conducted in this study, we observed that PSI substantially penalizes clustering results when the correct number of clusters is not recovered. In their work, Watson et al. adjusted the resolution hyperparameter to match the true number of clusters, or stopped the optimization after 1000 iterations if this was not achievable, thereby effectively reducing the extent to which PSI penalized the outcomes. Since we do not explicitly adjust algorithms to produce the correct number of clusters, we expect PSI performance to vary significantly.

#### 6.7 Wilcoxon signed-rank test results for comparing Leiden and Louvain algorithms

We assessed whether there were differences in performance between the two algorithms. We compared their performance distributions for the different clustering evaluation measures across the three scenarios (Max, Default, Mean). Using the Wilcoxon signed-rank test, we identified a statistical difference only in the mean performance scenario for the ARI and PSI measures, specifically in favor of Louvain. The Wilcoxon signed-rank test statistics (W) and corresponding p-values are reported in Table 4.

Table 4: P-values and test statistics for the Wilcoxon signed-rank test

| | ARI | | PSI | | NMI | |
|---|---|---|---|---|---|---|
| | p-value | W | p-value | W | p-value | W |
| Max | 0.50 | 5 | 0.89 | 7 | 0.28 | 1 |
| Default | 0.75 | 9 | 0.60 | 8 | 0.75 | 9 |
| Mean | **0.01** | 17 | **0.03** | 27 | 0.4 | 51 |

#### 6.8 Supplementary Figures and Tables

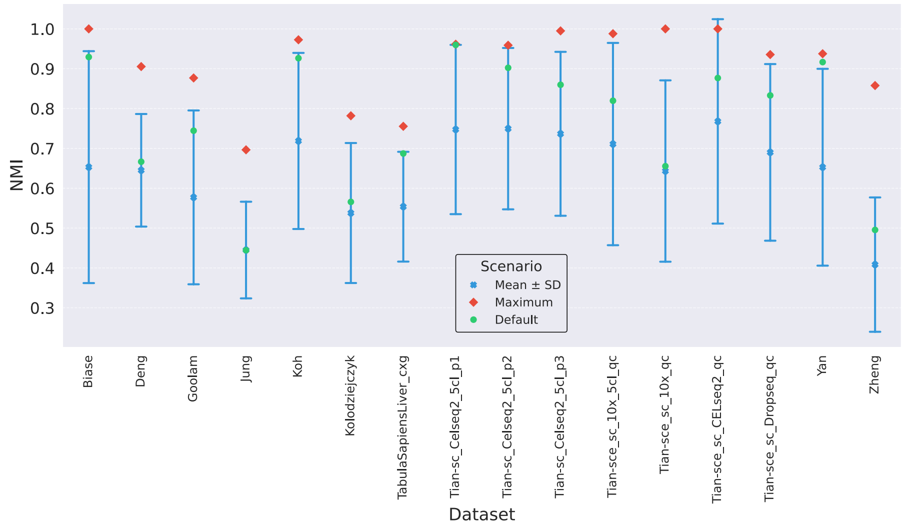


Figure 7: NMI scores per dataset for the Maximum, Default, and Mean performance scenarios for Leiden algorithm.

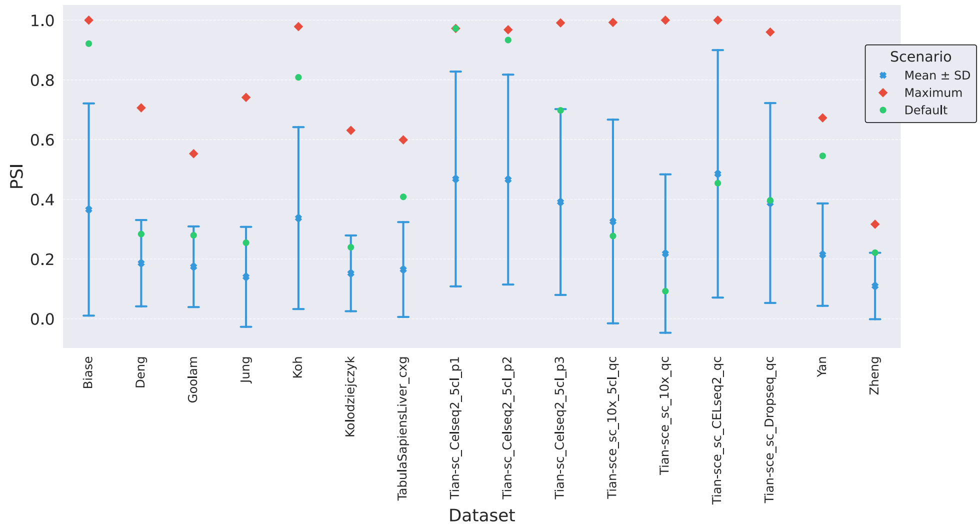


Figure 8: PSI scores per dataset for the Maximum, Default, and Mean performance scenario for Leiden algorithm.

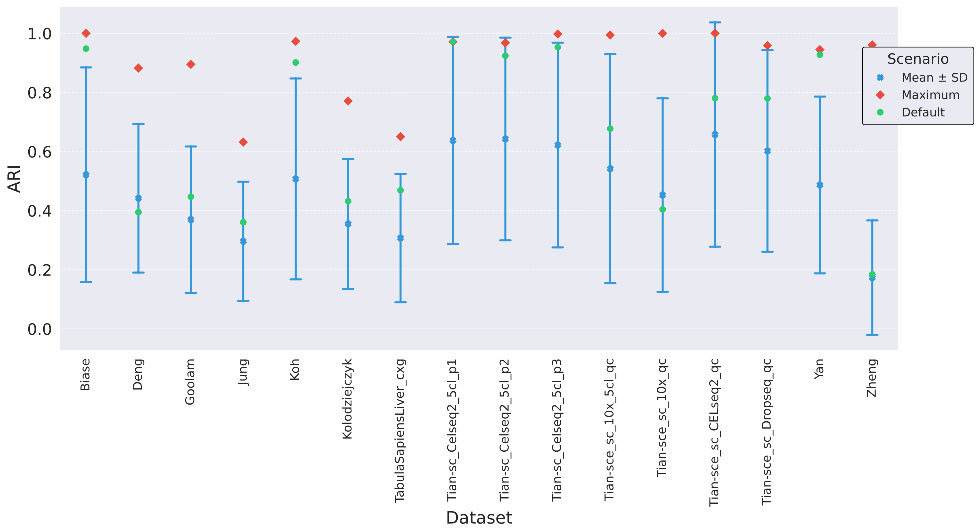


Figure 9: ARI scores per dataset for the Maximum, Default, and Mean performance scenarios for Louvain algorithm.

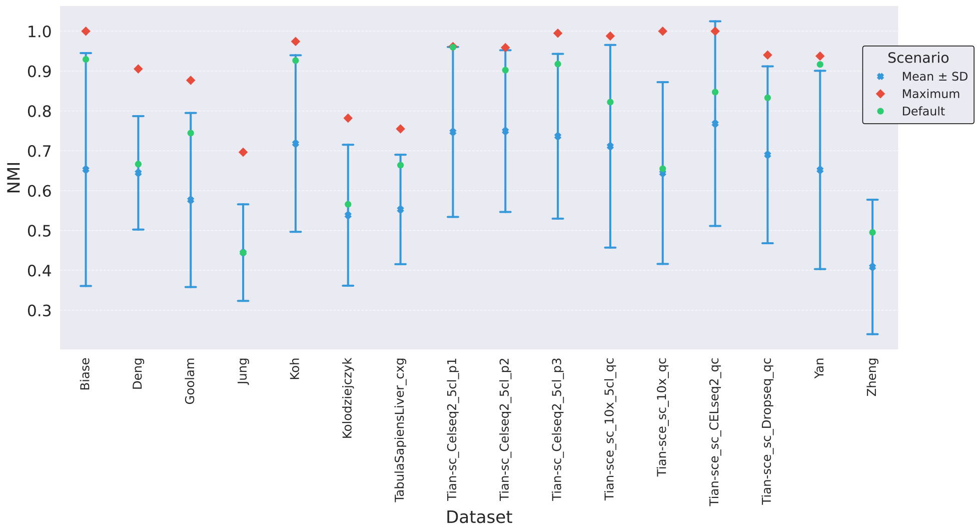


Figure 10: NMI scores per dataset for the Maximum, Default, and Mean performance scenario for Louvain algorithm.

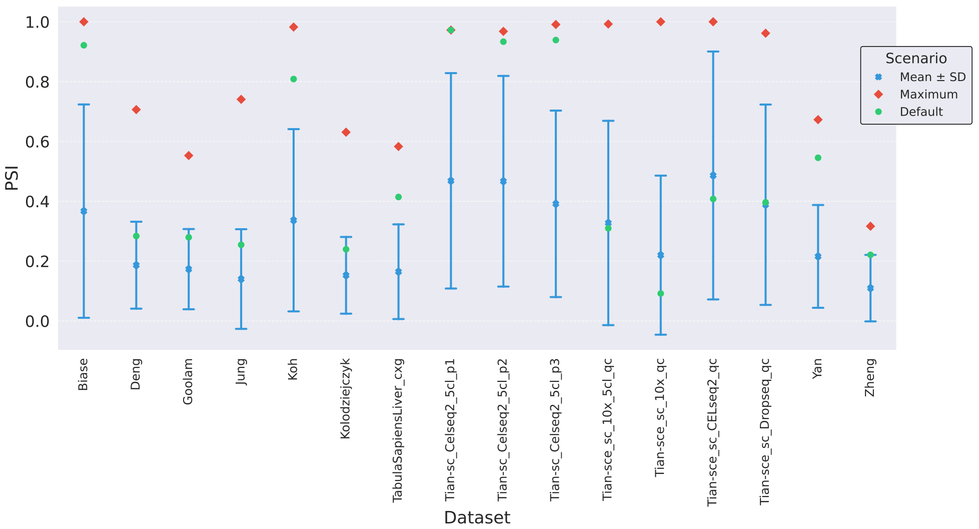


Figure 11: PSI scores per dataset for the Maximum, Default, and Mean performance scenario for Louvain algorithm.

Table 5: Best hyperparameter combinations for each dataset per algorithm. In many cases, we found multiple hyperparameter combinations achieving the same best ARI score. To resolve these ties, we first examined how many resolution values maintained this peak ARI score. Combinations that maintained the peak score across more resolution values were considered more robust and thus preferred. This approach effectively reduced the number of ties, but some still remained. As a second step, we compared the mean ARI across all resolution values for the tied hyperparameter combinations. Finally, this resolved all of the ties.

| Dataset | Clustering_algorithm | Quality_Function | Similarity_Measure | K_Neighbors | Graph_type | Resolution | ARI | NMI | PSI |
|---|---|---|---|---|---|---|---|---|---|
| Biase | leiden | RBERVertexPartition | phs | 14 | gauss | 1.161 | 1 | 1 | 1 |
| Biase | louvain | RBERVertexPartition | phs | 14 | gauss | 1.161 | 1 | 1 | 1 |
| Deng | louvain | RBERVertexPartition | braycurtis | 5 | umap | 0.201 | 0.882567177 | 0.905433929 | 0.706455152 |
| Deng | leiden | RBConfigurationVertexPartition | braycurtis | 5 | gauss | 0.201 | 0.882567177 | 0.905433929 | 0.706455152 |
| Goolam | louvain | CPMVertexPartition | cosine | 65 | jaccard_seurat | 1.401 | 0.894964327 | 0.852010823 | 0.505343971 |
| Goolam | leiden | CPMVertexPartition | cosine | 65 | jaccard_seurat | 1.401 | 0.894964327 | 0.852010823 | 0.505343971 |
| Jung | leiden | RBERVertexPartition | rho | 60 | umap | 0.941 | 0.644620071 | 0.677373012 | 0.44414406 |
| Jung | louvain | RBERVertexPartition | rho | 46 | umap | 0.861 | 0.63183291 | 0.672525488 | 0.53919184 |
| Koh | louvain | CPMVertexPartition | rho | 27 | gauss | 0.181 | 0.9731196 | 0.974287278 | 0.982401906 |
| Koh | leiden | CPMVertexPartition | phs | 32 | gauss | 0.241 | 0.968598137 | 0.971296596 | 0.978536131 |
| Kolodziejczyk | leiden | RBERVertexPartition | braycurtis | 39 | jaccard_phenograph | 0.301 | 0.771127153 | 0.778873818 | 0.339508617 |
| Kolodziejczyk | louvain | RBERVertexPartition | braycurtis | 37 | jaccard_phenograph | 0.381 | 0.771127153 | 0.778873818 | 0.339508617 |
| TabulaSapiensLiver_cxg | leiden | RBConfigurationVertexPartition | spearman | 52 | gauss | 1.341 | 0.642243081 | 0.732914558 | 0.364283044 |
| TabulaSapiensLiver_cxg | louvain | RBConfigurationVertexPartition | braycurtis | 11 | jaccard_phenograph | 0.021 | 0.650100626 | 0.704470557 | 0.065181385 |
| Tian-sc_Celseq2_5cl_p1 | louvain | RBConfigurationVertexPartition | euclidean | 20 | umap | 0.841 | 0.971283103 | 0.959749135 | 0.972306465 |
| Tian-sc_Celseq2_5cl_p1 | leiden | RBConfigurationVertexPartition | euclidean | 20 | umap | 0.841 | 0.971283103 | 0.959749135 | 0.972306465 |
| Tian-sc_Celseq2_5cl_p2 | leiden | RBERVertexPartition | euclidean | 40 | gauss | 1.001 | 0.967867795 | 0.947463742 | 0.965232632 |
| Tian-sc_Celseq2_5cl_p2 | louvain | RBERVertexPartition | euclidean | 40 | gauss | 1.001 | 0.967867795 | 0.947463742 | 0.965232632 |
| Tian-sc_Celseq2_5cl_p3 | louvain | RBERVertexPartition | spearman | 25 | gauss | 0.901 | 0.997999993 | 0.994973348 | 0.794579147 |
| Tian-sc_Celseq2_5cl_p3 | leiden | RBERVertexPartition | spearman | 25 | gauss | 0.901 | 0.997999993 | 0.994973348 | 0.794579147 |
| Tian-sce_sc_10x_5cl_qc | leiden | RBConfigurationVertexPartition | correlation | 30 | jaccard_seurat | 0.101 | 0.99421975 | 0.98791548 | 0.992307885 |
| Tian-sce_sc_10x_5cl_qc | louvain | RBConfigurationVertexPartition | correlation | 30 | jaccard_seurat | 0.101 | 0.99421975 | 0.98791548 | 0.992307885 |
| Tian-sce_sc_10x_qc | leiden | RBConfigurationVertexPartition | braycurtis | 100 | gauss | 0.281 | 1 | 1 | 1 |
| Tian-sce_sc_10x_qc | louvain | RBConfigurationVertexPartition | braycurtis | 100 | gauss | 0.281 | 1 | 1 | 1 |
| Tian-sce_sc_CELseq2_qc | louvain | RBConfigurationVertexPartition | braycurtis | 51 | jaccard_seurat | 1.021 | 1 | 1 | 1 |
| Tian-sce_sc_CELseq2_qc | leiden | RBConfigurationVertexPartition | braycurtis | 51 | jaccard_seurat | 1.021 | 1 | 1 | 1 |
| Tian-sce_sc_Dropseq_qc | louvain | RBConfigurationVertexPartition | cosine | 29 | gauss | 0.281 | 0.958769585 | 0.940312604 | 0.961765354 |
| Tian-sce_sc_Dropseq_qc | leiden | RBConfigurationVertexPartition | euclidean | 36 | jaccard_seurat | 1.321 | 0.947029177 | 0.918518327 | 0.637580808 |
| Yan | leiden | RBERVertexPartition | cosine | 21 | gauss | 1.601 | 0.945040753 | 0.937603616 | 0.567307692 |
| Yan | louvain | RBERVertexPartition | cosine | 21 | gauss | 1.681 | 0.945040753 | 0.937603616 | 0.567307692 |
| Zheng | leiden | RBConfigurationVertexPartition | braycurtis | 7 | gauss | 0.021 | 0.959567436 | 0.857790896 | 0.282988879 |
| Zheng | louvain | RBConfigurationVertexPartition | correlation | 7 | umap | 0.021 | 0.960760011 | 0.854302405 | 0.285195215 |

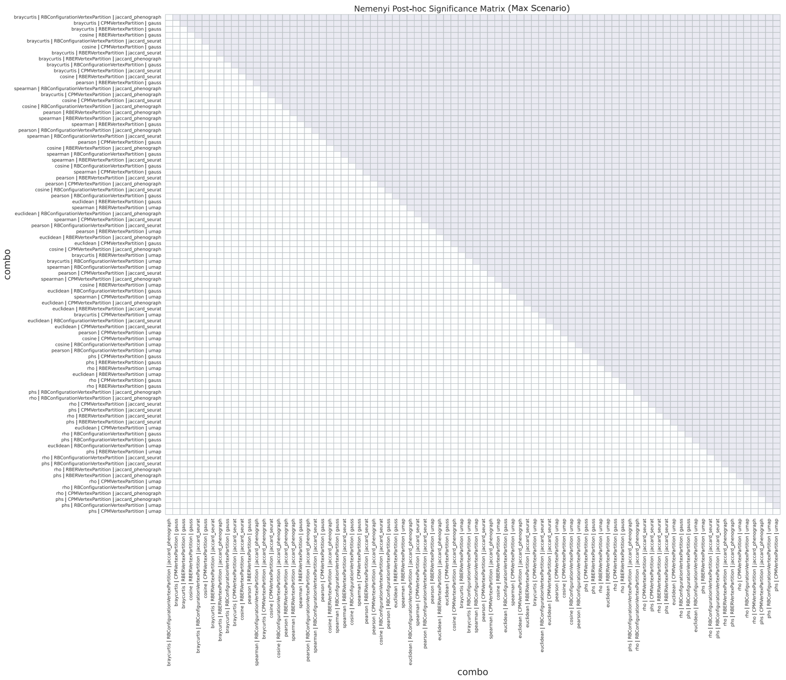


(a) Maximum Scenario for Leiden ($p = 7.33 \times 10^{-12}$). No critical difference was found when performance was maximized.

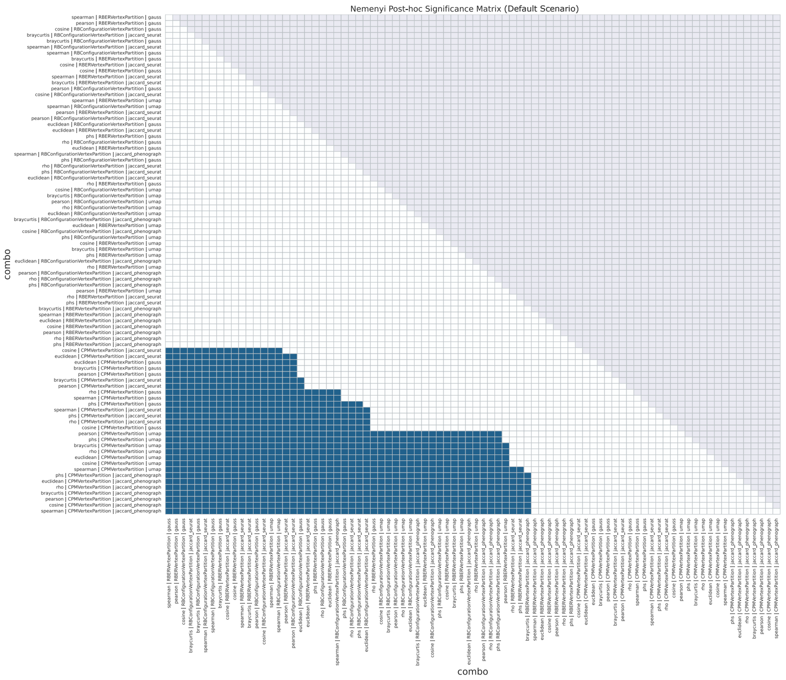


(b) Default Scenario for Leiden ($p = 1.20 \times 10^{-146}$)

Figure 12: Pairwise statistical significance matrices for the Leiden algorithm (Part I)

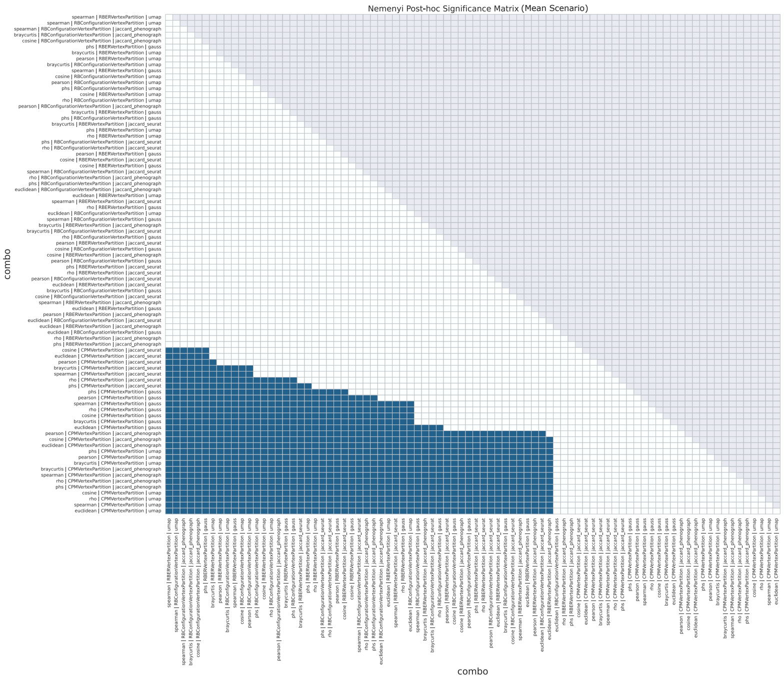


(c) Mean Scenario for Leiden ($p = 4.63 \times 10^{-138}$)

Figure 12: Pairwise statistical significance matrices for the Leiden algorithm (Part II). The binary heatmaps represent significant pairwise differences ($p < 0.05$) derived from the Nemenyi post-hoc test. The test was applied following a Friedman test which confirmed global significance in all scenarios. However, the Nemenyi test didn't discover any critically different combinations in the maximum scenario. Both the x and y axes represent the 84 hyperparameter combinations, sorted from the best (top-left) to the worst (bottom-right) according to their average rank.

Table 6: Clustering Scores Comparison

| | ARI_mean | | NMI_mean | | PSI_mean | | ARI_default | | NMI_default | | PSI_default | | ARI_max | | NMI_max | | PSI_max | |
|---|---|---|---|---|---|---|---|---|---|---|---|---|---|---|---|---|---|---|
| Clustering_algorithm | leiden | louvain | leiden | louvain | leiden | louvain | leiden | louvain | leiden | louvain | leiden | louvain | leiden | louvain | leiden | louvain | leiden | louvain |
| Biase | 0.5205 | **0.5212** | **0.6530** | **0.6530** | 0.3658 | **0.3671** | **0.9483** | **0.9483** | **0.9293** | **0.9293** | **0.9214** | **0.9214** | **1.0000** | **1.0000** | **1.0000** | **1.0000** | **1.0000** | **1.0000** |
| Deng | 0.4415 | **0.4418** | **0.6451** | 0.6447 | 0.1861 | **0.1864** | **0.3949** | **0.3949** | **0.6666** | **0.6666** | **0.2837** | **0.2837** | **0.8826** | **0.8826** | **0.9054** | **0.9054** | **0.7065** | **0.7065** |
| Goolam | 0.3693 | **0.3694** | **0.5771** | 0.5765 | **0.1741** | 0.1731 | **0.4472** | **0.4472** | **0.7444** | **0.7444** | **0.2795** | **0.2795** | **0.8950** | **0.8950** | **0.8768** | **0.8768** | **0.5529** | **0.5529** |
| Jung | **0.2964** | **0.2964** | **0.4449** | 0.4446 | **0.1403** | 0.1400 | **0.3606** | **0.3606** | **0.4445** | **0.4445** | **0.2545** | **0.2545** | **0.6446** | 0.6318 | **0.6965** | **0.6965** | **0.7413** | 0.7407 |
| Koh | **0.5079** | 0.5074 | **0.7187** | 0.7182 | **0.3371** | 0.3366 | **0.9014** | **0.9014** | **0.9264** | **0.9264** | **0.8086** | **0.8086** | 0.9686 | **0.9731** | 0.9726 | **0.9743** | 0.9785 | **0.9824** |
| Kolodziejczyk | 0.3537 | **0.3551** | 0.5378 | **0.5384** | 0.1521 | **0.1525** | **0.4318** | **0.4318** | **0.5658** | **0.5658** | **0.2395** | **0.2395** | **0.7711** | **0.7711** | **0.7819** | **0.7819** | **0.6309** | **0.6309** |
| TabulaSapiensLiver_cxg | **0.3076** | 0.3071 | **0.5537** | 0.5528 | **0.1648** | 0.1646 | **0.4829** | 0.4690 | **0.6872** | 0.6639 | 0.4082 | **0.4143** | 0.6422 | **0.6501** | **0.7554** | 0.7550 | **0.5992** | 0.5831 |
| Tian-sc_Celseq2_5cl_p1 | 0.6375 | **0.6376** | **0.7474** | 0.7472 | 0.4681 | **0.4684** | **0.9713** | **0.9713** | **0.9597** | **0.9597** | **0.9723** | **0.9723** | **0.9713** | **0.9713** | **0.9614** | **0.9614** | **0.9723** | **0.9723** |
| Tian-sc_Celseq2_5cl_p2 | 0.6420 | **0.6425** | **0.7496** | 0.7494 | 0.4661 | **0.4669** | **0.9241** | **0.9241** | **0.9023** | **0.9023** | **0.9335** | **0.9335** | **0.9679** | **0.9679** | **0.9589** | **0.9589** | **0.9680** | **0.9680** |
| Tian-sc_Celseq2_5cl_p3 | 0.6218 | **0.6220** | **0.7366** | 0.7364 | 0.3909 | **0.3915** | 0.7731 | **0.9529** | 0.8597 | **0.9177** | 0.6979 | **0.9389** | **0.9980** | **0.9980** | **0.9950** | **0.9950** | **0.9910** | **0.9910** |
| Tian-sce_sc_10x_5cl_qc | 0.5406 | **0.5417** | 0.7109 | **0.7113** | 0.3258 | **0.3275** | 0.6724 | **0.6775** | 0.8196 | **0.8222** | 0.2772 | **0.3097** | **0.9942** | **0.9942** | **0.9879** | **0.9879** | **0.9924** | **0.9924** |
| Tian-sce_sc_10x_qc | 0.4508 | **0.4527** | 0.6432 | **0.6442** | 0.2183 | **0.2199** | **0.4060** | 0.4044 | **0.6554** | 0.6550 | **0.0924** | 0.0917 | **1.0000** | **1.0000** | **1.0000** | **1.0000** | **1.0000** | **1.0000** |
| Tian-sce_sc_CELseq2_qc | 0.6566 | **0.6575** | 0.7678 | **0.7682** | 0.4853 | **0.4862** | **0.8588** | 0.7804 | **0.8767** | 0.8472 | **0.4541** | 0.4078 | **1.0000** | **1.0000** | **1.0000** | **1.0000** | **1.0000** | **1.0000** |
| Tian-sce_sc_Dropseq_qc | 0.6018 | **0.6021** | **0.6900** | **0.6900** | 0.3876 | **0.3885** | **0.7796** | **0.7796** | **0.8330** | **0.8330** | **0.3965** | **0.3965** | 0.9470 | **0.9588** | 0.9356 | **0.9403** | 0.9602 | **0.9618** |
| Yan | 0.4855 | **0.4868** | **0.6527** | 0.6520 | 0.2148 | **0.2157** | **0.9277** | **0.9277** | **0.9166** | **0.9166** | **0.5454** | **0.5454** | **0.9450** | **0.9450** | **0.9376** | **0.9376** | **0.6729** | **0.6729** |
| Zheng | 0.1725 | **0.1730** | 0.4084 | **0.4086** | **0.1097** | 0.1096 | **0.1852** | 0.1842 | **0.4955** | 0.4952 | **0.2212** | 0.2210 | 0.9596 | **0.9608** | **0.8578** | **0.8578** | 0.3165 | **0.3166** |

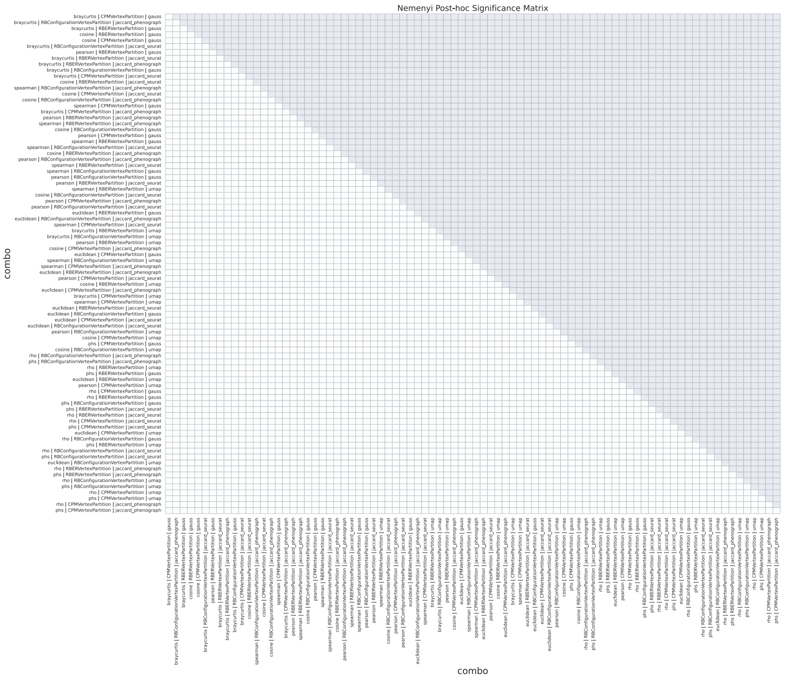


(a) Maximum Scenario for Louvain ($p = 4.88 \times 10^{-13}$)

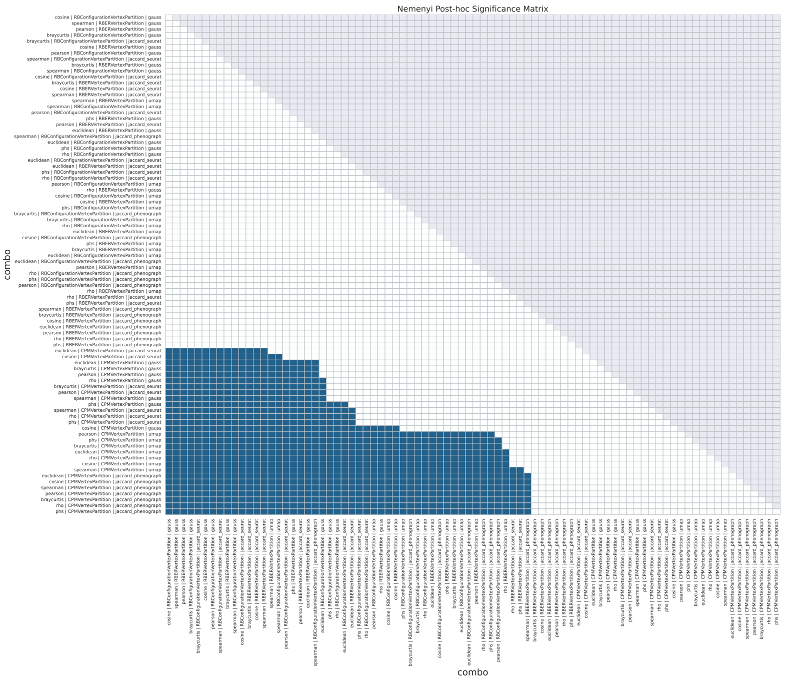


(b) Default Scenario for Louvain ($p = 5.46 \times 10^{-147}$)

Figure 13: Pairwise statistical significance matrices for the Louvain algorithm (Part I)

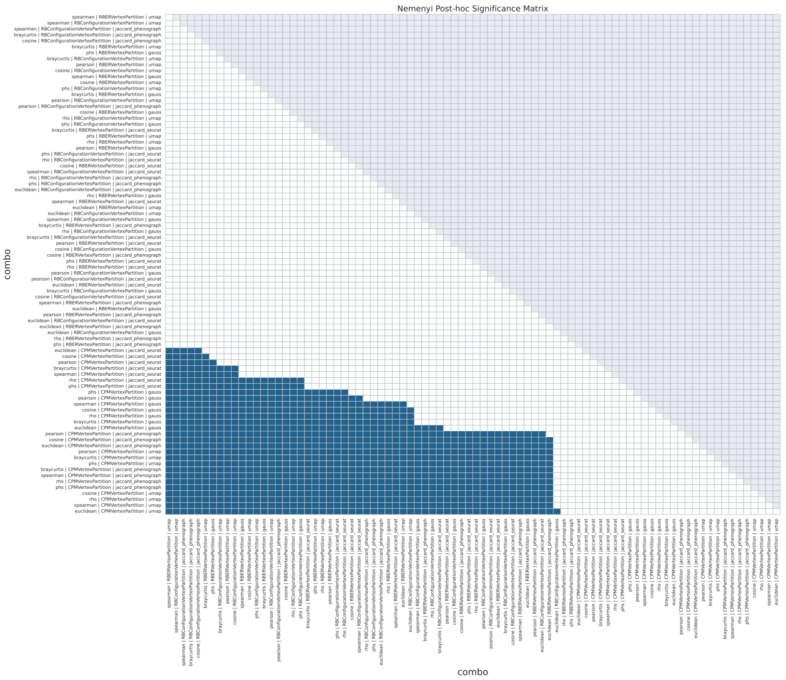


(c) Mean Scenario for Louvain ($p = 3.28 \times 10^{-138}$)

Figure 13: Pairwise statistical significance matrices for the Louvain algorithm (Part II). The binary heatmaps represent significant pairwise differences ($p < 0.05$) derived from the Nemenyi post-hoc test. The test was applied following a Friedman test which confirmed global significance in all scenarios. However, the Nemenyi test didn't discover any critically different combinations in the maximum scenario. Both the x and y axes represent the 84 hyperparameter combinations, sorted from the best (top-left) to the worst (bottom-right) according to their average rank.

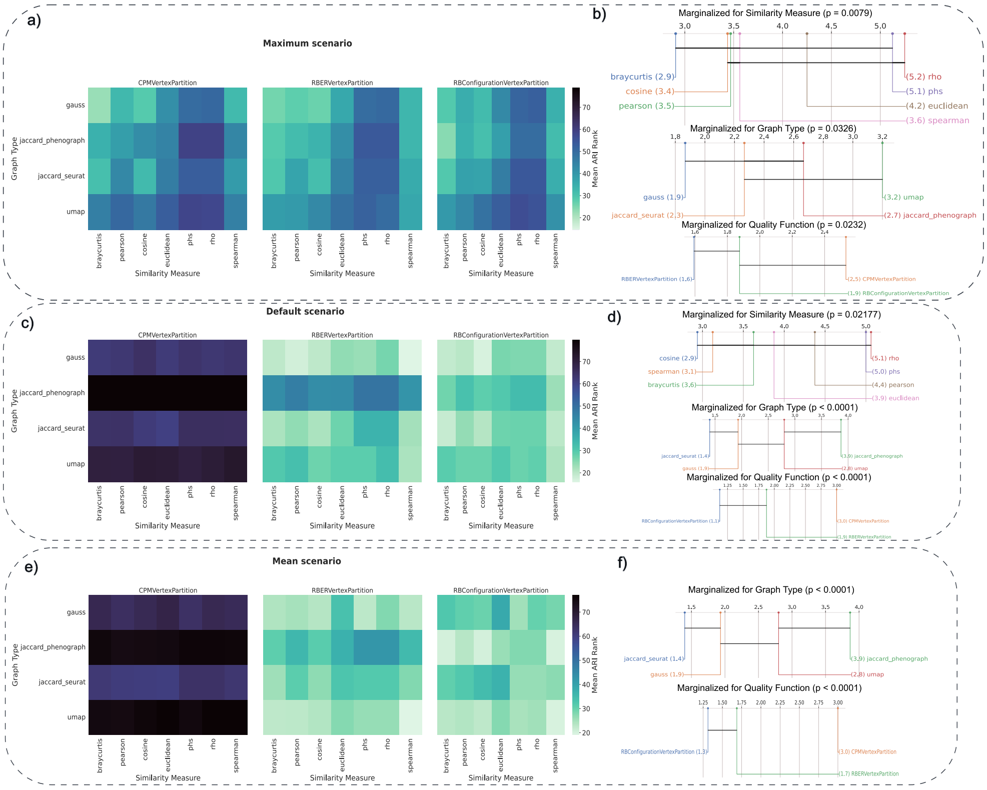


Figure 14: Performance landscapes illustrating the interaction between numerical ($k$, Resolution) and categorical hyperparameters across representative datasets for the Louvain algorithm. Each column corresponds to a dataset (Zheng, Jung, Koh, Tian-sc_Celseq2_5cl_p1), and each row represents a different scenario: Maximum configuration (top), Default configuration (middle), and Maximum mean configuration (bottom). Color intensity reflects ARI scores, where bright regions indicate high clustering performance and dark regions indicate poor clustering results. The pink circle marks the optimal numerical coordinates identified in the Maximum configuration, while the blue triangle indicates the default Scanpy settings.

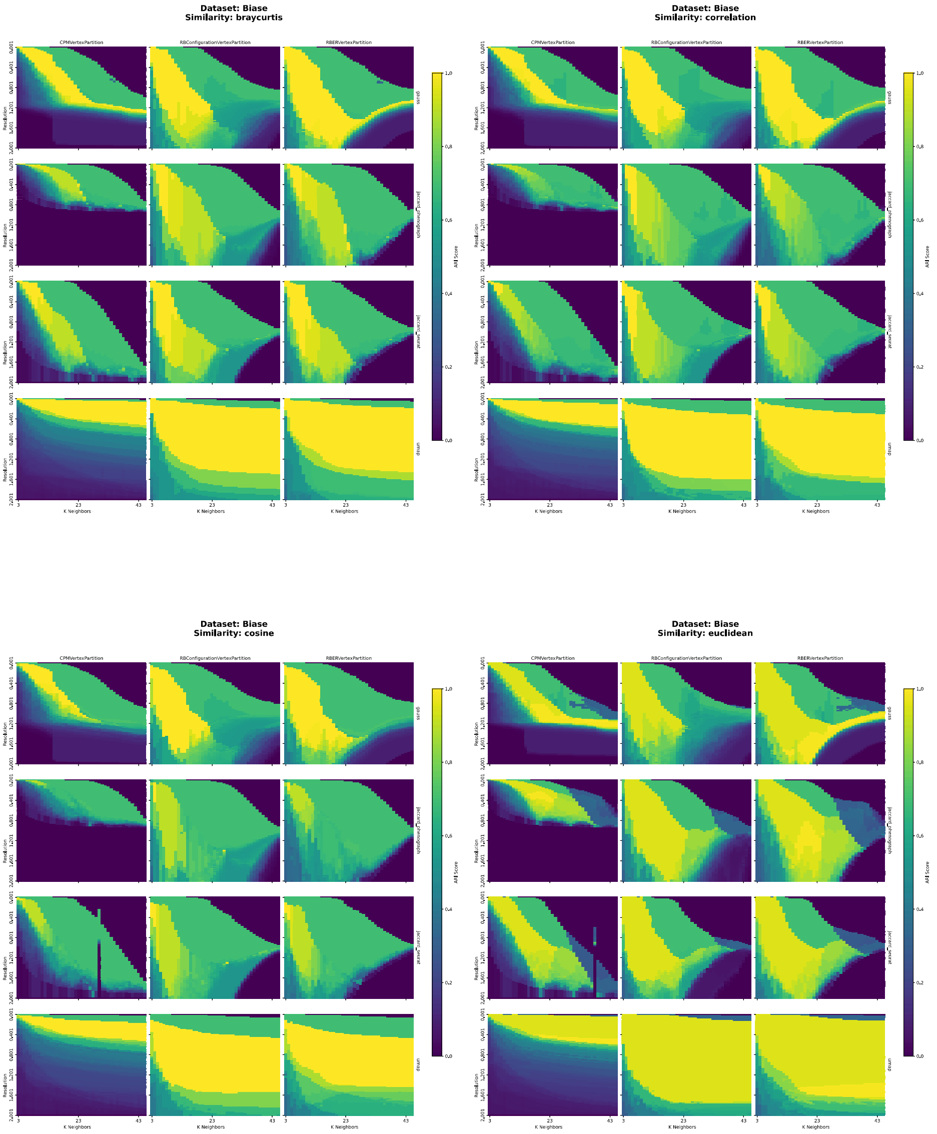


Figure 15: Performance landscapes for Biase, for: `braycurtis`, `correlation`, `euclidean`, `cosine`. Performance landscapes illustrate the interaction between numerical hyperparameters ($k$, resolution) and categorical hyperparameters. Each 12-cell grid represents combinations of Graph Type and Quality Function for a given Similarity Measure. Color intensity encodes ARI scores, with brighter regions indicating higher clustering performance and darker regions indicating poorer results.

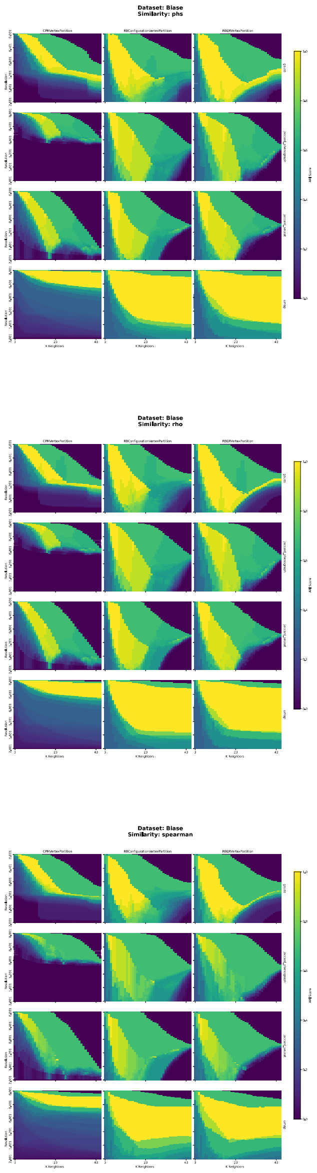


Figure 16: Performance landscapes for Biase, for: `phs`, `rho`, `spearman`. Performance landscapes illustrate the interaction between numerical hyperparameters ($k$, resolution) and categorical hyperparameters. Each 12-cell grid represents combinations of Graph Type and Quality Function for a given Similarity Measure. Color intensity encodes ARI scores, with brighter regions indicating higher clustering performance and darker regions indicating poorer results.

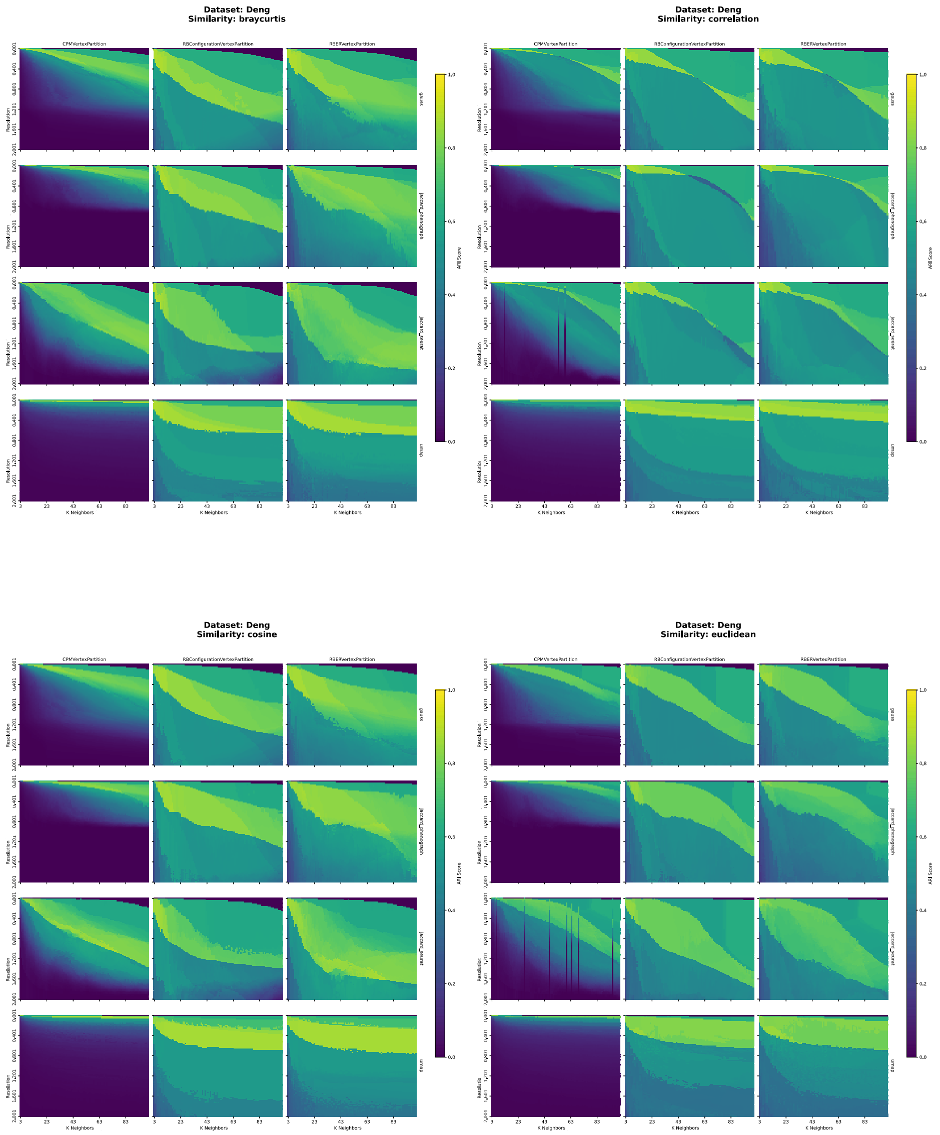


Figure 17: Performance landscapes for Deng, for: `braycurtis`, `correlation`, `euclidean`, `cosine`. Performance landscapes illustrate the interaction between numerical hyperparameters ($k$, resolution) and categorical hyperparameters. Each 12-cell grid represents combinations of Graph Type and Quality Function for a given Similarity Measure. Color intensity encodes ARI scores, with brighter regions indicating higher clustering performance and darker regions indicating poorer results.

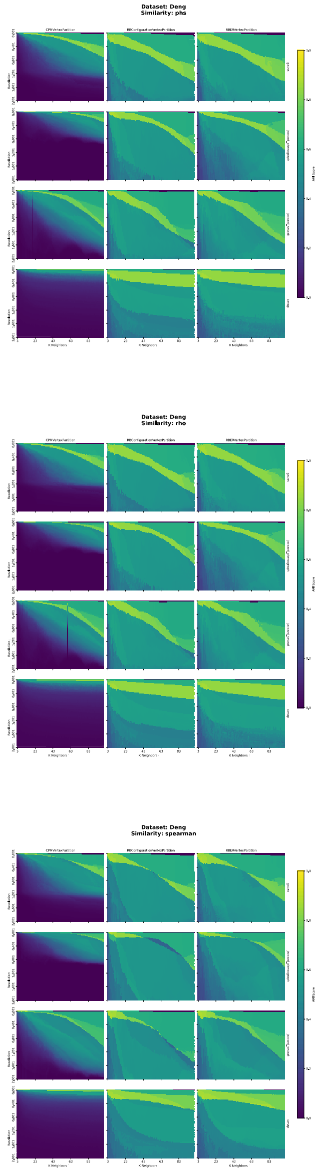


Figure 18: Performance landscapes for Deng, for: `phs`, `rho`, `spearman`. Performance landscapes illustrate the interaction between numerical hyperparameters ($k$, resolution) and categorical hyperparameters. Each 12-cell grid represents combinations of Graph Type and Quality Function for a given Similarity Measure. Color intensity encodes ARI scores, with brighter regions indicating higher clustering performance and darker regions indicating poorer results.

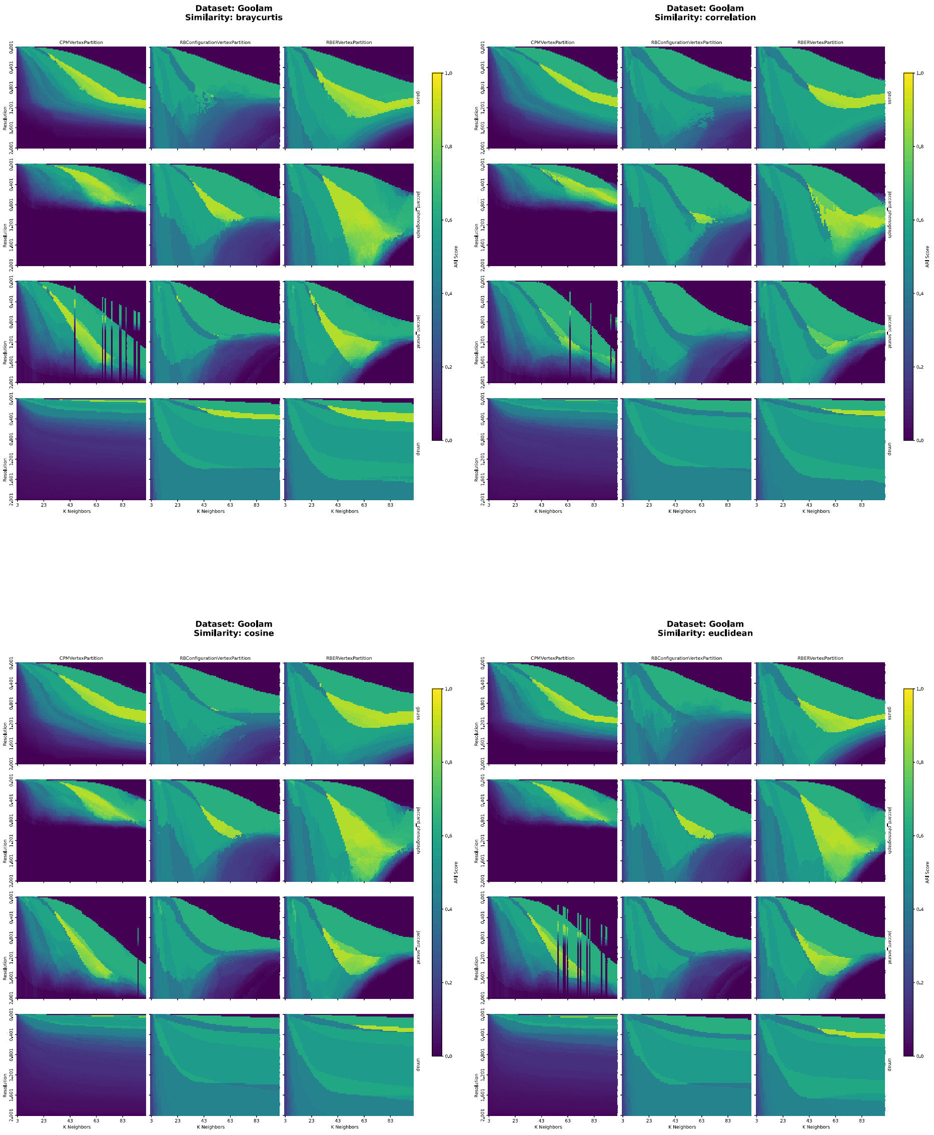


Figure 19: Performance landscapes for Goolam, for: `braycurtis`, `correlation`, `euclidean`, `cosine`. Performance landscapes illustrate the interaction between numerical hyperparameters ($k$, resolution) and categorical hyperparameters. Each 12-cell grid represents combinations of Graph Type and Quality Function for a given Similarity Measure. Color intensity encodes ARI scores, with brighter regions indicating higher clustering performance and darker regions indicating poorer results.

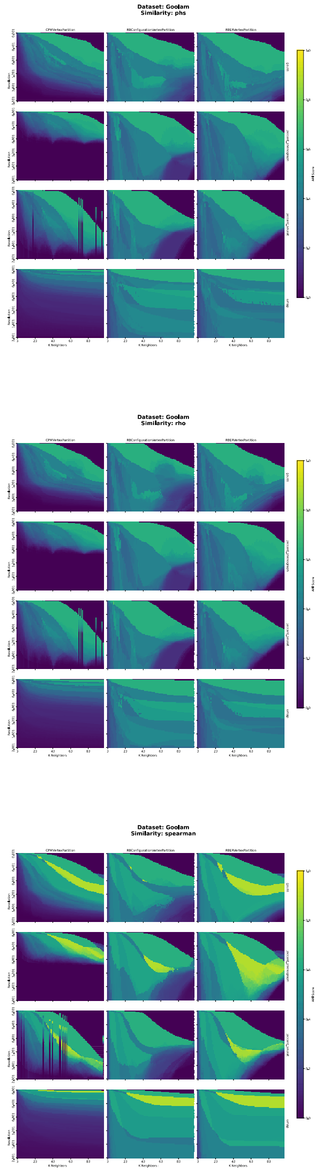


Figure 20: Performance landscapes for Goolam, for: `phs`, `rho`, `spearman`. Performance landscapes illustrate the interaction between numerical hyperparameters ($k$, resolution) and categorical hyperparameters. Each 12-cell grid represents combinations of Graph Type and Quality Function for a given Similarity Measure. Color intensity encodes ARI scores, with brighter regions indicating higher clustering performance and darker regions indicating poorer results.

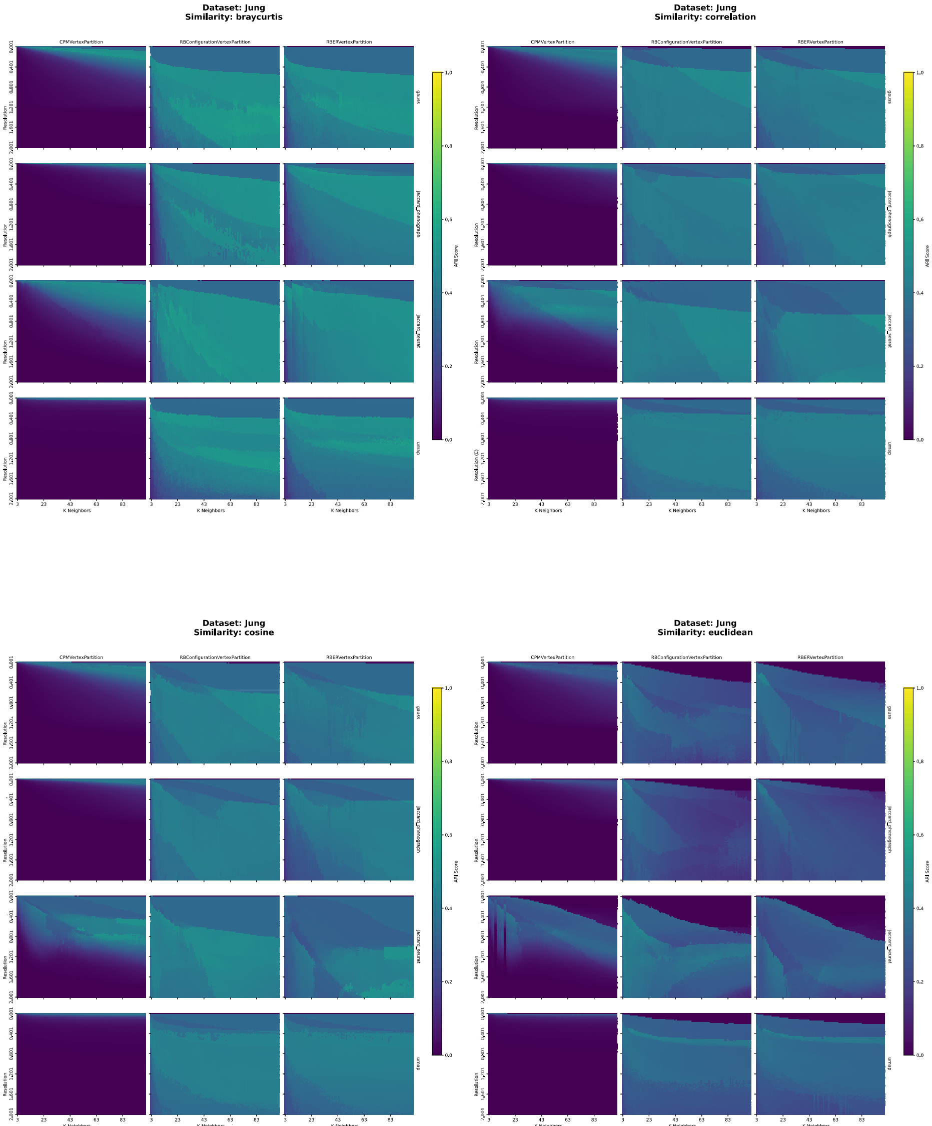


Figure 21: Performance landscapes for Jung, for: `braycurtis`, `correlation`, `euclidean`, `cosine`. Performance landscapes illustrate the interaction between numerical hyperparameters ($k$, resolution) and categorical hyperparameters. Each 12-cell grid represents combinations of Graph Type and Quality Function for a given Similarity Measure. Color intensity encodes ARI scores, with brighter regions indicating higher clustering performance and darker regions indicating poorer results.

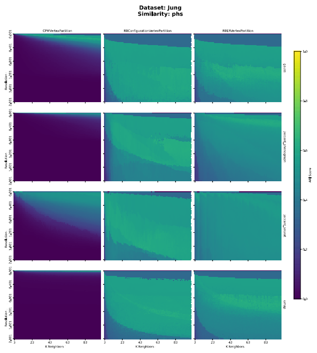


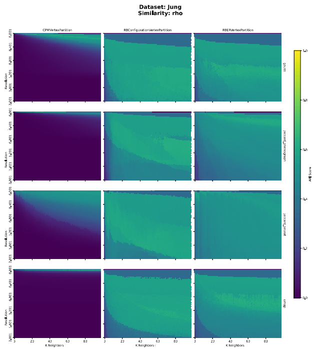


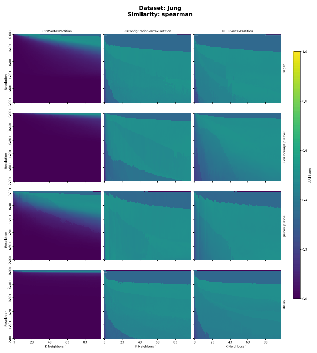


Figure 22: Performance landscapes for Jung, for: `phs`, `rho`, `spearman`. Performance landscapes illustrate the interaction between numerical hyperparameters ($k$, resolution) and categorical hyperparameters. Each 12-cell grid represents combinations of Graph Type and Quality Function for a given Similarity Measure. Color intensity encodes ARI scores, with brighter regions indicating higher clustering performance and darker regions indicating poorer results.

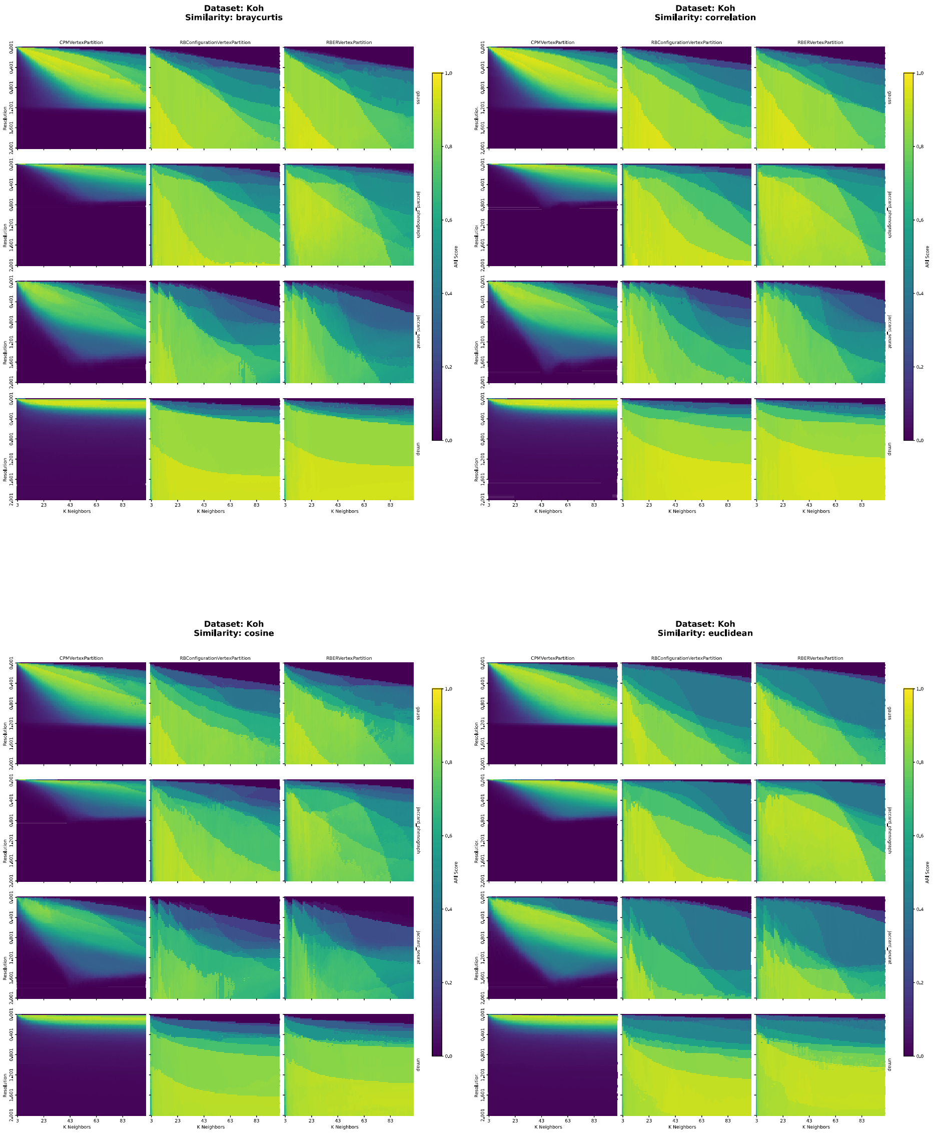


Figure 23: Performance landscapes for Koh, for: `braycurtis`, `correlation`, `euclidean`, `cosine`. Performance landscapes illustrate the interaction between numerical hyperparameters ($k$, resolution) and categorical hyperparameters. Each 12-cell grid represents combinations of Graph Type and Quality Function for a given Similarity Measure. Color intensity encodes ARI scores, with brighter regions indicating higher clustering performance and darker regions indicating poorer results.

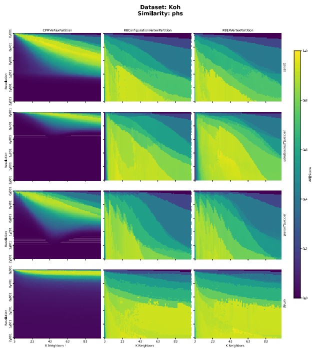


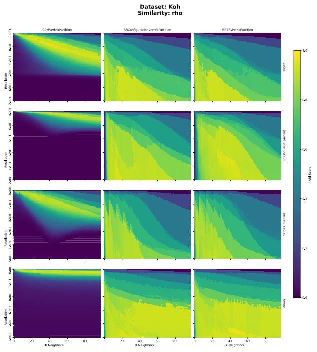


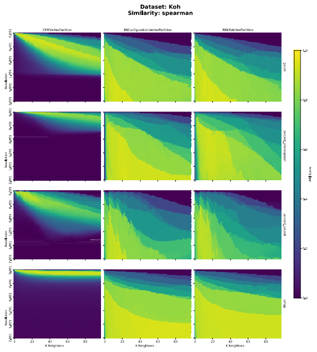


Figure 24: Performance landscapes for Koh, for: `phs`, `rho`, `spearman`. Performance landscapes illustrate the interaction between numerical hyperparameters ($k$, resolution) and categorical hyperparameters. Each 12-cell grid represents combinations of Graph Type and Quality Function for a given Similarity Measure. Color intensity encodes ARI scores, with brighter regions indicating higher clustering performance and darker regions indicating poorer results.

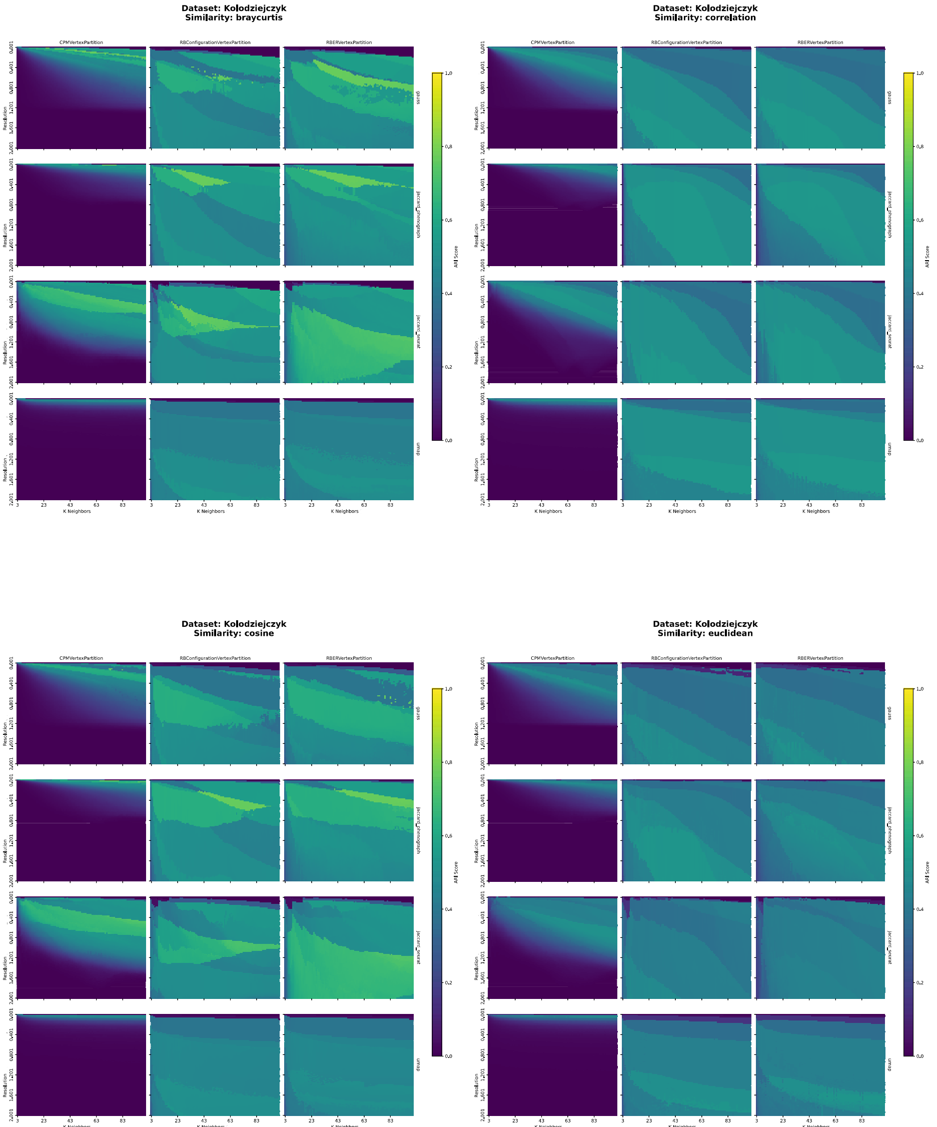


Figure 25: Performance landscapes for Kolodziejczyk, for: `braycurtis`, `correlation`, `euclidean`, `cosine`. Performance landscapes illustrate the interaction between numerical hyperparameters ($k$, resolution) and categorical hyperparameters. Each 12-cell grid represents combinations of Graph Type and Quality Function for a given Similarity Measure. Color intensity encodes ARI scores, with brighter regions indicating higher clustering performance and darker regions indicating poorer results.

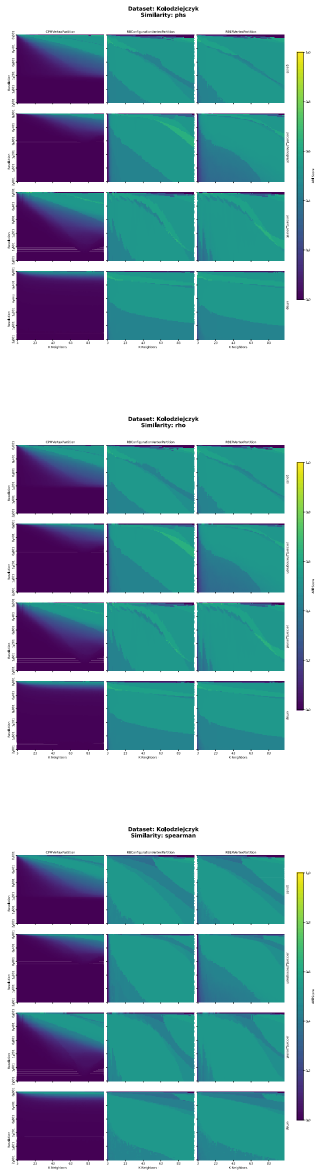


Figure 26: Performance landscapes for Kolodziejczyk, for: `phs`, `rho`, `spearman`. Performance landscapes illustrate the interaction between numerical hyperparameters ($k$, resolution) and categorical hyperparameters. Each 12-cell grid represents combinations of Graph Type and Quality Function for a given Similarity Measure. Color intensity encodes ARI scores, with brighter regions indicating higher clustering performance and darker regions indicating poorer results.

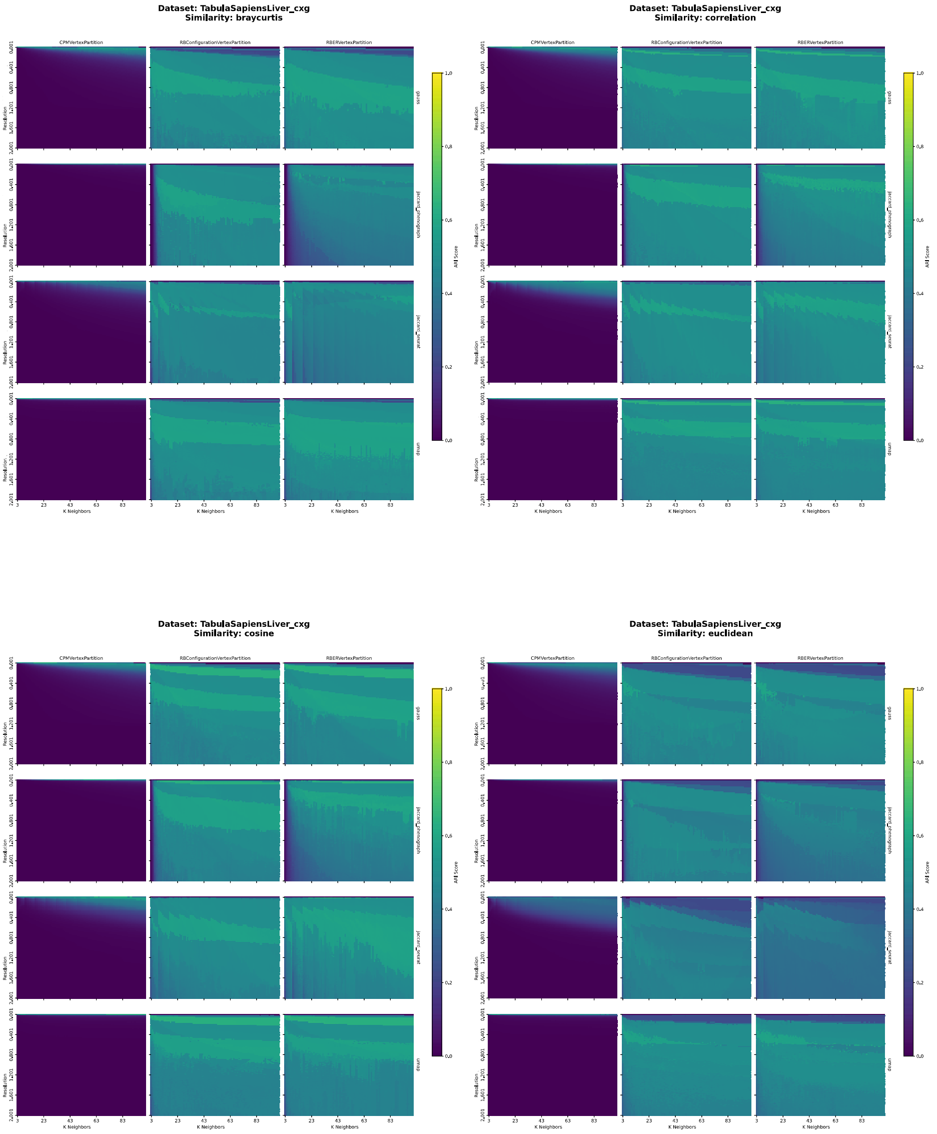


Figure 27: Performance landscapes for TabulaSapiensLiver_cxg, for: `braycurtis`, `correlation`, `euclidean`, `cosine`. Performance landscapes illustrate the interaction between numerical hyperparameters ($k$, resolution) and categorical hyperparameters. Each 12-cell grid represents combinations of Graph Type and Quality Function for a given Similarity Measure. Color intensity encodes ARI scores, with brighter regions indicating higher clustering performance and darker regions indicating poorer results.

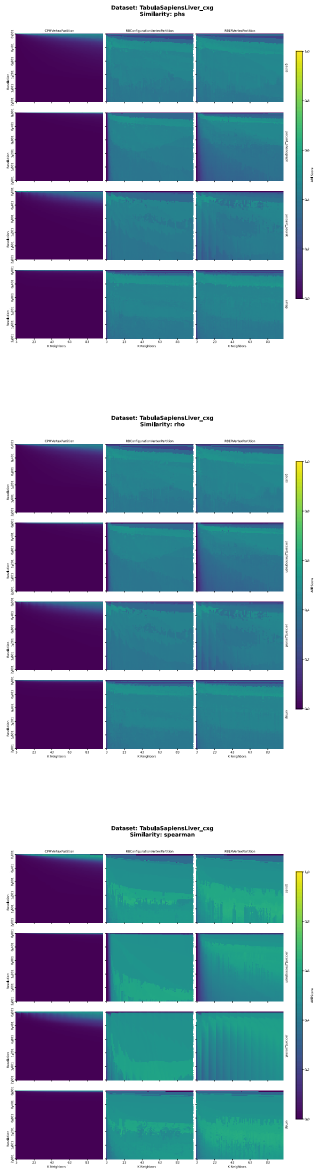


Figure 28: Performance landscapes for TabulaSapiensLiver_cxg, for: `phs`, `rho`, `spearman`. Performance landscapes illustrate the interaction between numerical hyperparameters ($k$, resolution) and categorical hyperparameters. Each 12-cell grid represents combinations of Graph Type and Quality Function for a given Similarity Measure. Color intensity encodes ARI scores, with brighter regions indicating higher clustering performance and darker regions indicating poorer results.

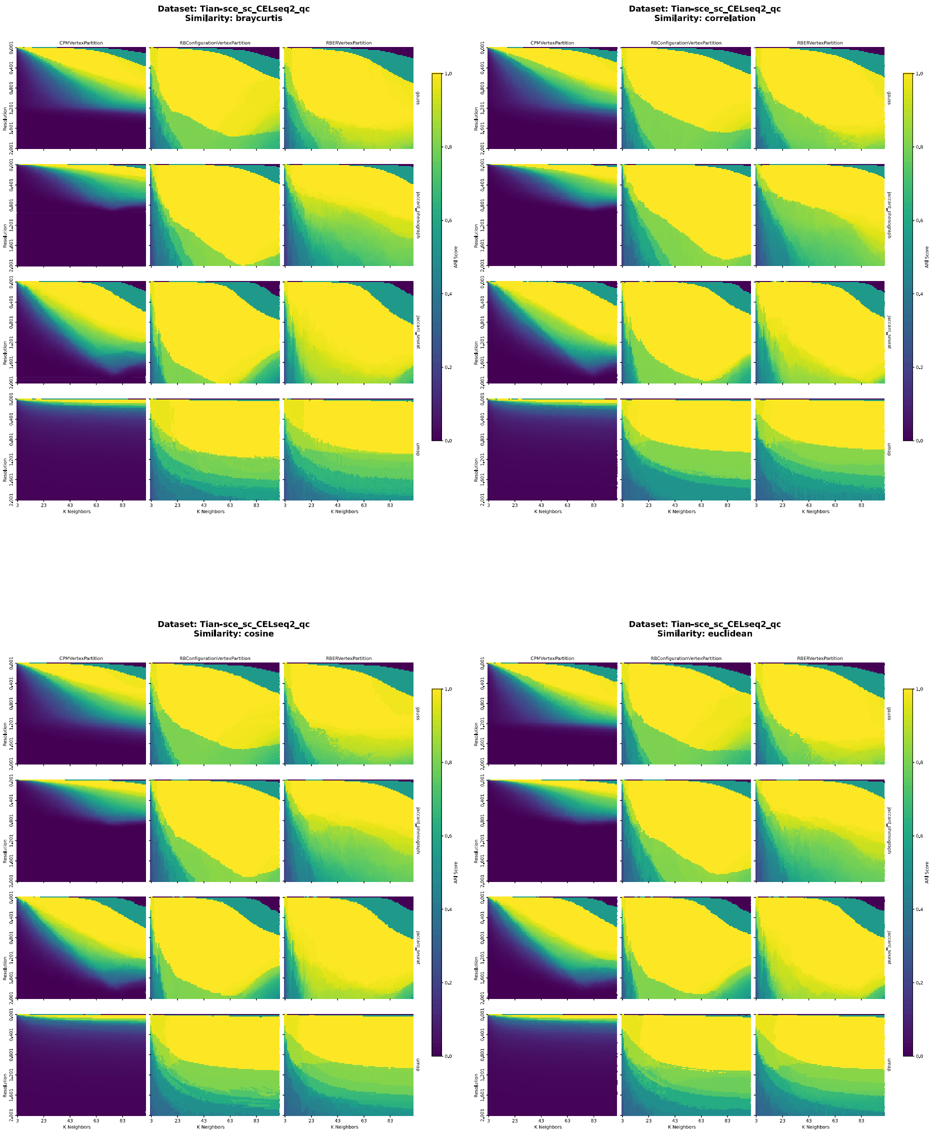


Figure 29: Performance landscapes for Tian-sce_sc_CELseq2_qc, for: `braycurtis`, `correlation`, `euclidean`, `cosine`. Performance landscapes illustrate the interaction between numerical hyperparameters ($k$, resolution) and categorical hyperparameters. Each 12-cell grid represents combinations of Graph Type and Quality Function for a given Similarity Measure. Color intensity encodes ARI scores, with brighter regions indicating higher clustering performance and darker regions indicating poorer results.

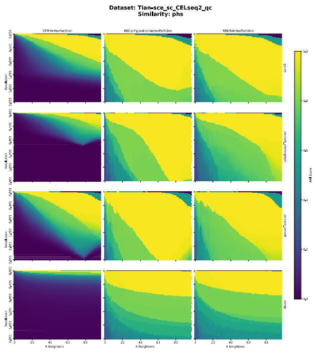


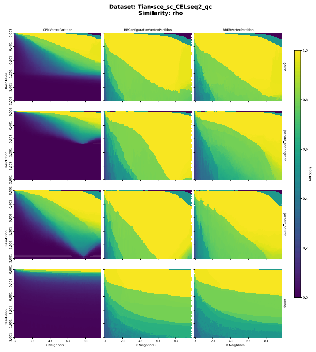


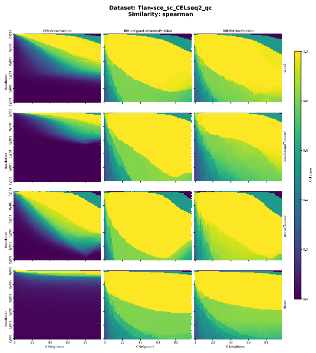


Figure 30: Performance landscapes for Tian-sce_sc_CELseq2_qc, for: `phs`, `rho`, `spearman`. Performance landscapes illustrate the interaction between numerical hyperparameters ($k$, resolution) and categorical hyperparameters. Each 12-cell grid represents combinations of Graph Type and Quality Function for a given Similarity Measure. Color intensity encodes ARI scores, with brighter regions indicating higher clustering performance and darker regions indicating poorer results.

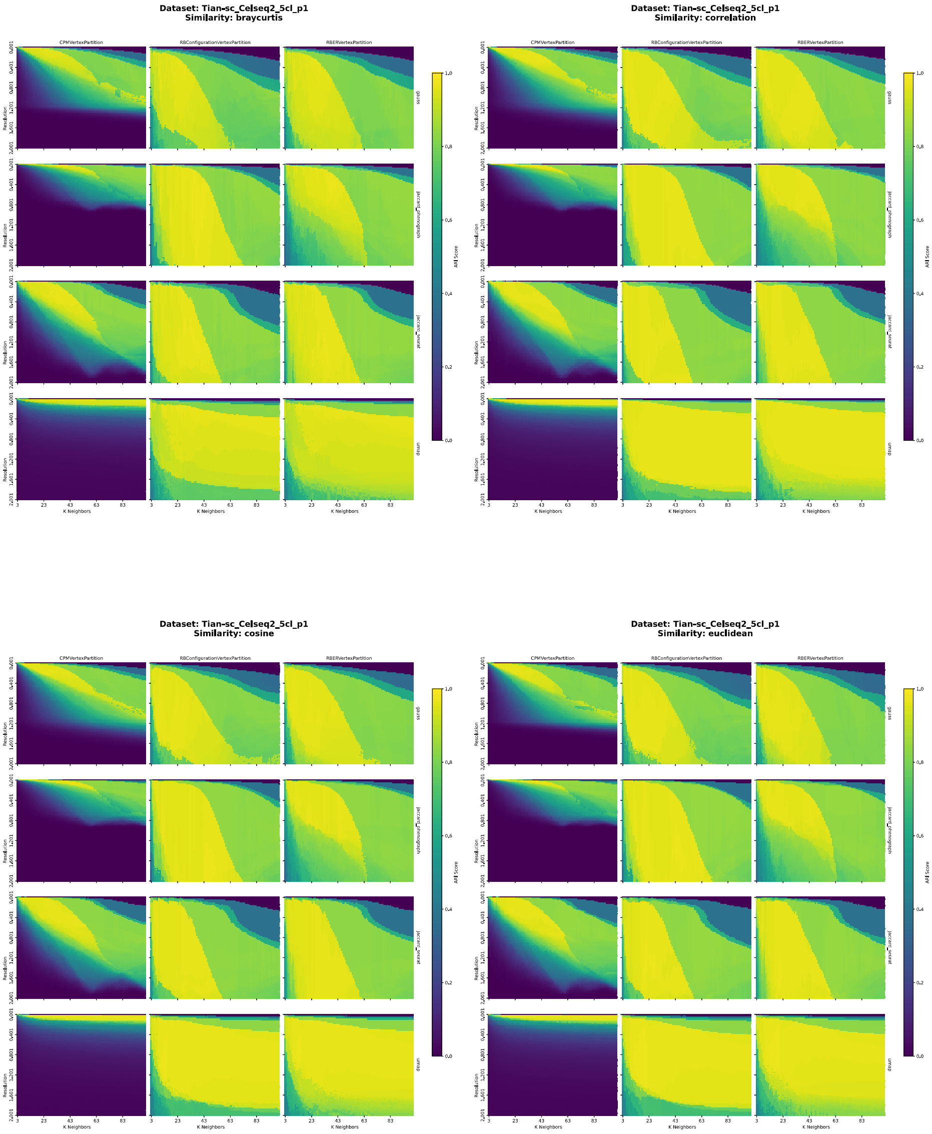


Figure 31: Performance landscapes for Tian-sc_Celseq2_5cl_p1, for: `braycurtis`, `correlation`, `euclidean`, `cosine`. Performance landscapes illustrate the interaction between numerical hyperparameters ($k$, resolution) and categorical hyperparameters. Each 12-cell grid represents combinations of Graph Type and Quality Function for a given Similarity Measure. Color intensity encodes ARI scores, with brighter regions indicating higher clustering performance and darker regions indicating poorer results.

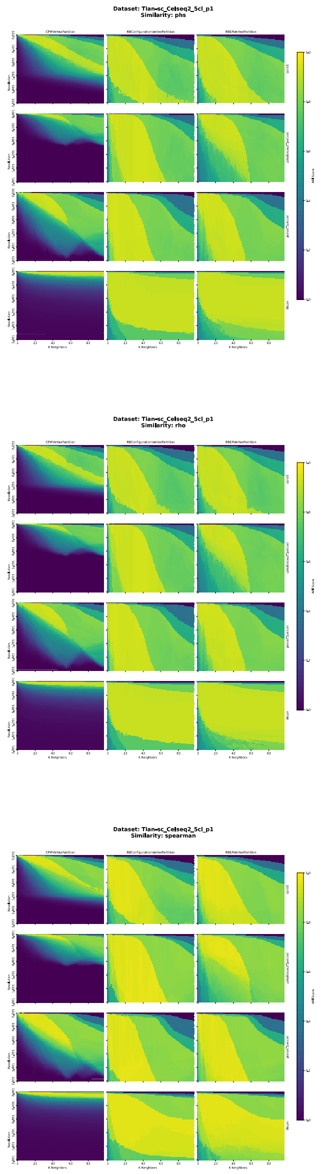


Figure 32: Performance landscapes for Tian-sc_Celseq2_5cl_p1, for: `phs`, `rho`, `spearman`. Performance landscapes illustrate the interaction between numerical hyperparameters ($k$, resolution) and categorical hyperparameters. Each 12-cell grid represents combinations of Graph Type and Quality Function for a given Similarity Measure. Color intensity encodes ARI scores, with brighter regions indicating higher clustering performance and darker regions indicating poorer results.

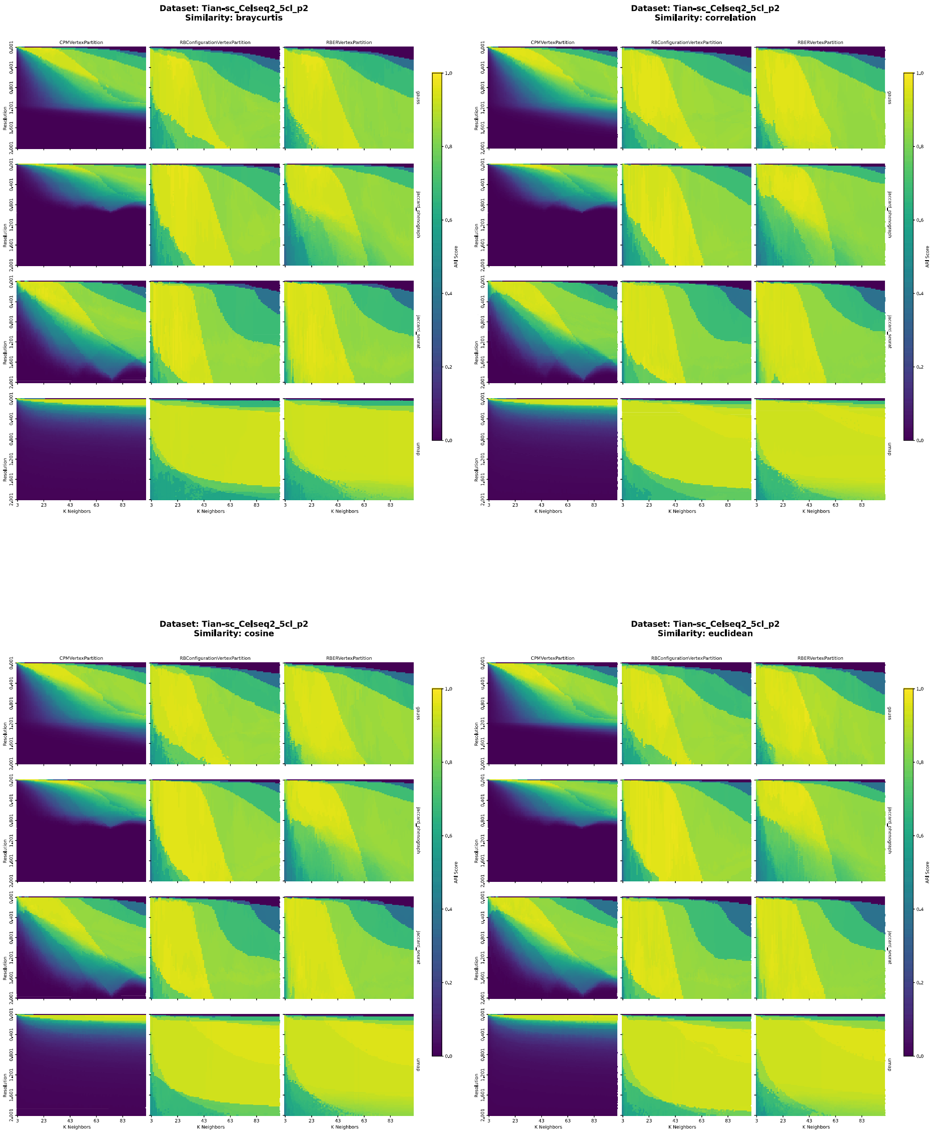


Figure 33: Performance landscapes for Tian-sc_Celseq2_5cl_p2, for: `braycurtis`, `correlation`, `euclidean`, `cosine`. Performance landscapes illustrate the interaction between numerical hyperparameters ($k$, resolution) and categorical hyperparameters. Each 12-cell grid represents combinations of Graph Type and Quality Function for a given Similarity Measure. Color intensity encodes ARI scores, with brighter regions indicating higher clustering performance and darker regions indicating poorer results.

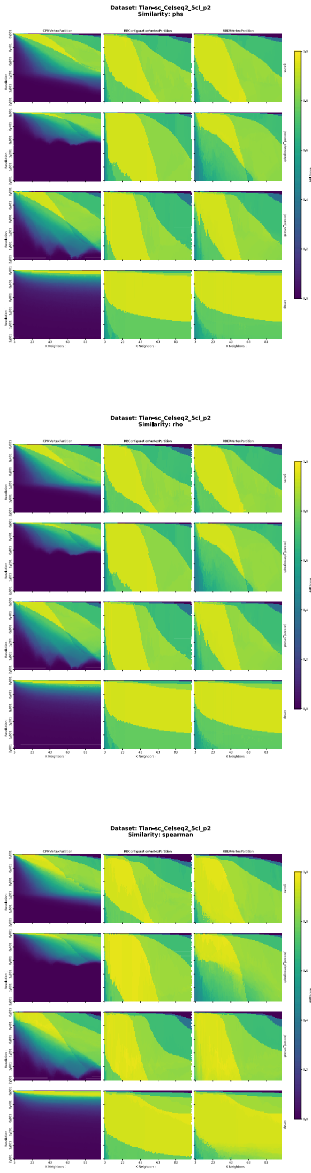


Figure 34: Performance landscapes for Tian-sc_Celseq2_5cl_p2, for: `phs`, `rho`, `spearman`. Performance landscapes illustrate the interaction between numerical hyperparameters ($k$, resolution) and categorical hyperparameters. Each 12-cell grid represents combinations of Graph Type and Quality Function for a given Similarity Measure. Color intensity encodes ARI scores, with brighter regions indicating higher clustering performance and darker regions indicating poorer results.

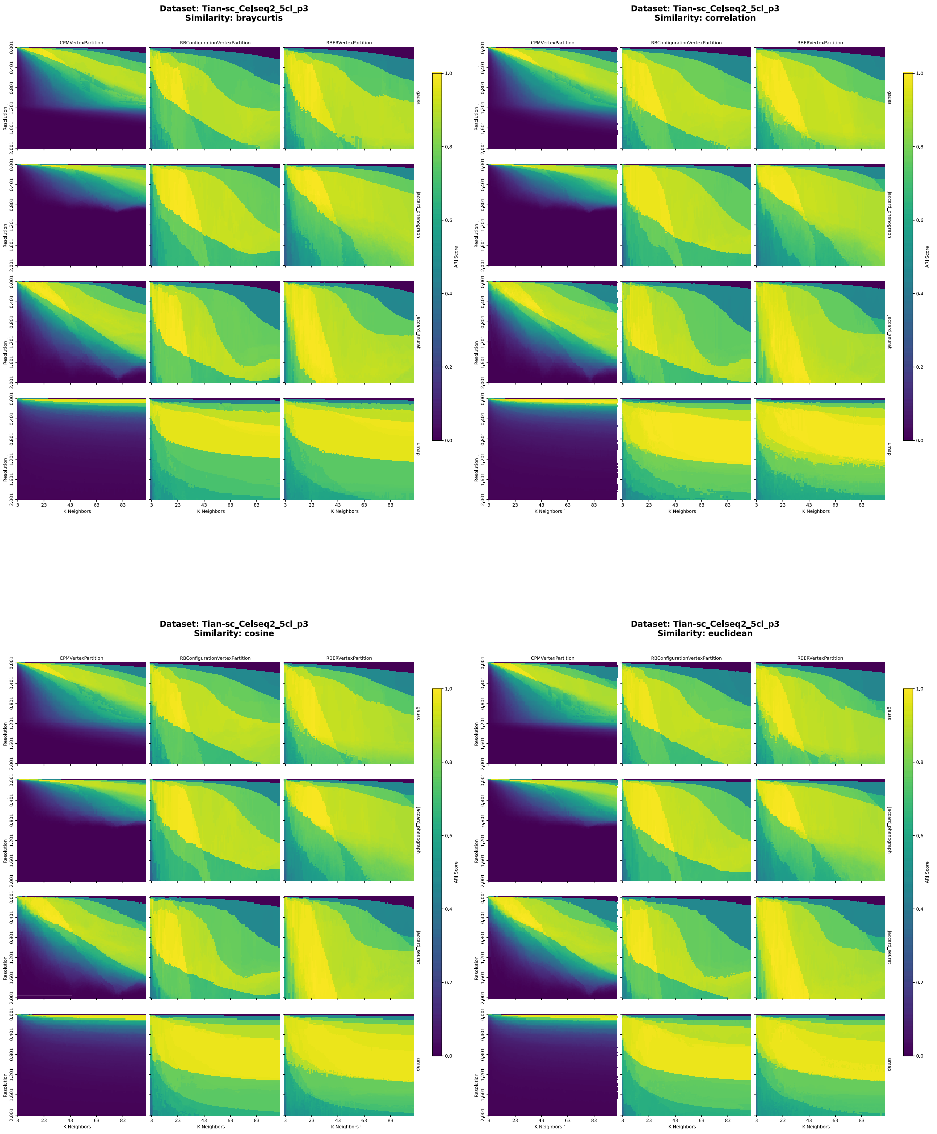


Figure 35: Performance landscapes for Tian-sc_Celseq2_5cl_p3, for: `braycurtis`, `correlation`, `euclidean`, `cosine`. Performance landscapes illustrate the interaction between numerical hyperparameters ($k$, resolution) and categorical hyperparameters. Each 12-cell grid represents combinations of Graph Type and Quality Function for a given Similarity Measure. Color intensity encodes ARI scores, with brighter regions indicating higher clustering performance and darker regions indicating poorer results.

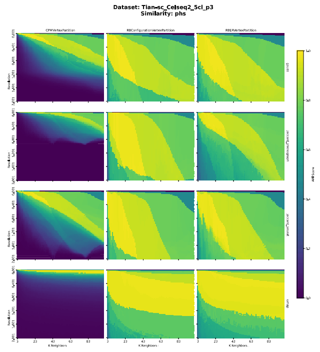


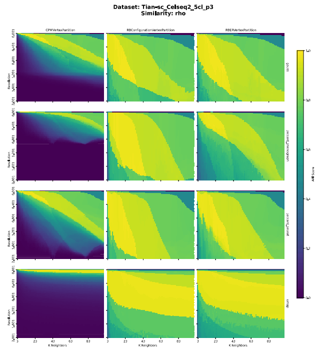


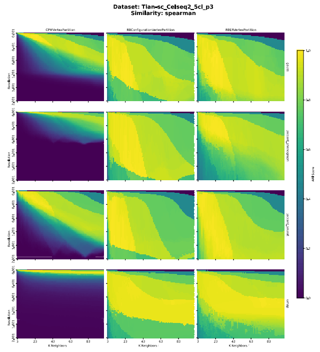


Figure 36: Performance landscapes for Tian-sc_Celseq2_5cl_p3, for: `phs`, `rho`, `spearman`. Performance landscapes illustrate the interaction between numerical hyperparameters ($k$, resolution) and categorical hyperparameters. Each 12-cell grid represents combinations of Graph Type and Quality Function for a given Similarity Measure. Color intensity encodes ARI scores, with brighter regions indicating higher clustering performance and darker regions indicating poorer results.

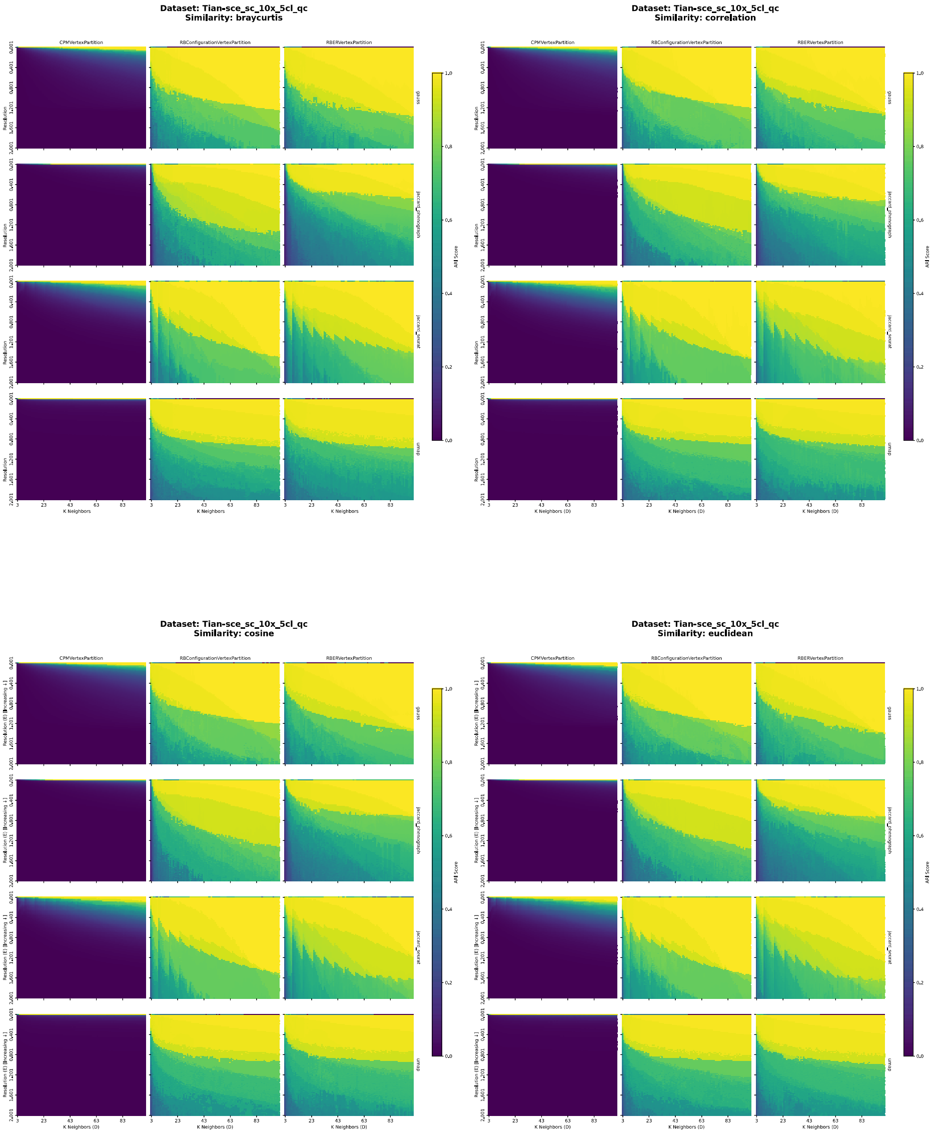


Figure 37: Performance landscapes for Tian-sce_sc_10x_5cl_qc, for: `braycurtis`, `correlation`, `euclidean`, `cosine`. Performance landscapes illustrate the interaction between numerical hyperparameters ($k$, resolution) and categorical hyperparameters. Each 12-cell grid represents combinations of Graph Type and Quality Function for a given Similarity Measure. Color intensity encodes ARI scores, with brighter regions indicating higher clustering performance and darker regions indicating poorer results.

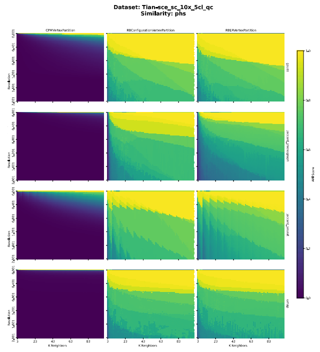


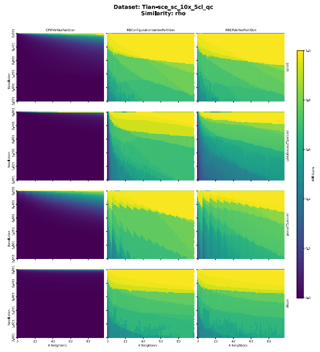


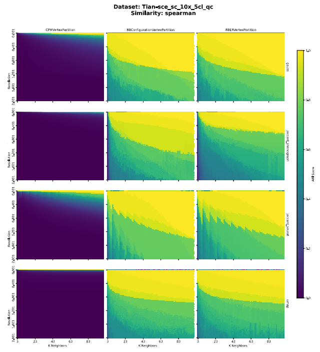


Figure 38: Performance landscapes for Tian-sce_sc_10x_5cl_qc, for: `phs`, `rho`, `spearman`. Performance landscapes illustrate the interaction between numerical hyperparameters ($k$, resolution) and categorical hyperparameters. Each 12-cell grid represents combinations of Graph Type and Quality Function for a given Similarity Measure. Color intensity encodes ARI scores, with brighter regions indicating higher clustering performance and darker regions indicating poorer results.

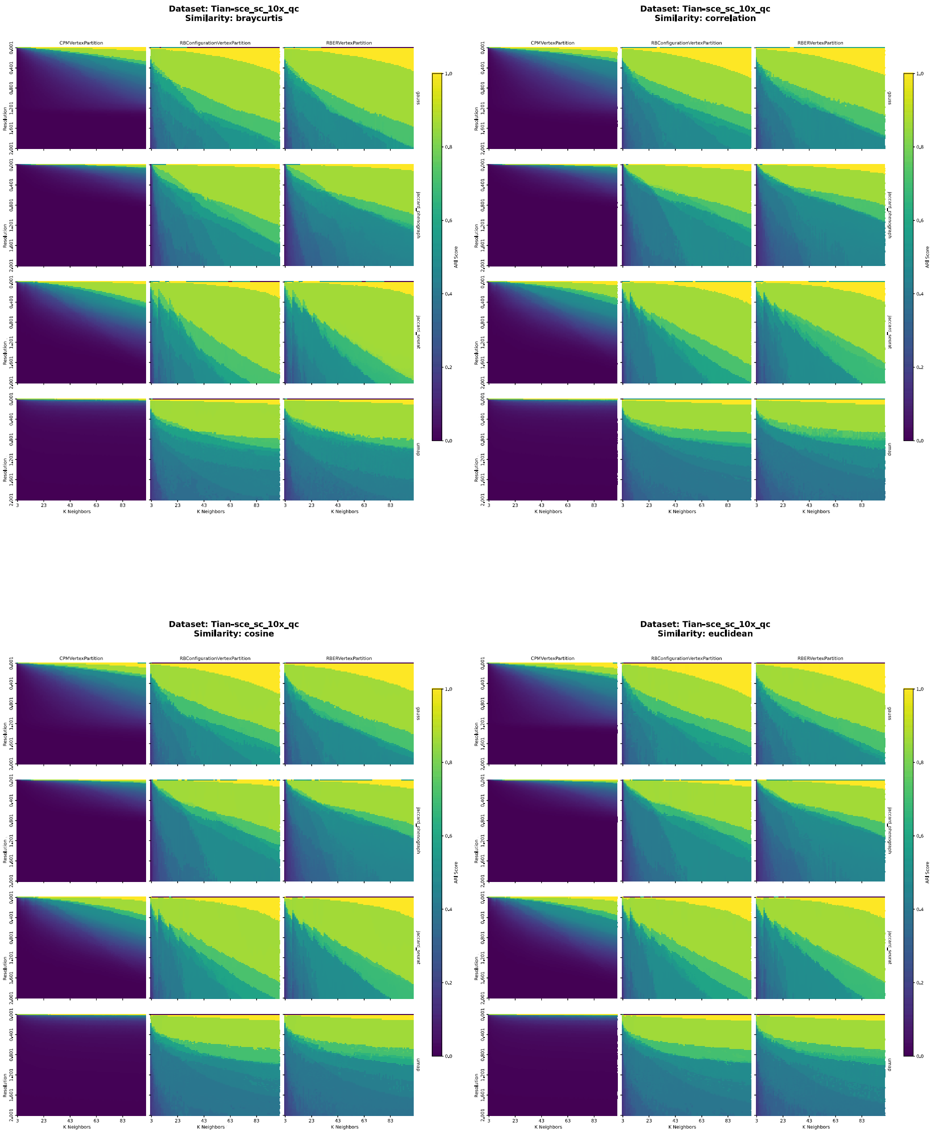


Figure 39: Performance landscapes for Tian-sce_sc_10x_qc, for: `braycurtis`, `correlation`, `euclidean`, `cosine`. Performance landscapes illustrate the interaction between numerical hyperparameters ($k$, resolution) and categorical hyperparameters. Each 12-cell grid represents combinations of Graph Type and Quality Function for a given Similarity Measure. Color intensity encodes ARI scores, with brighter regions indicating higher clustering performance and darker regions indicating poorer results.

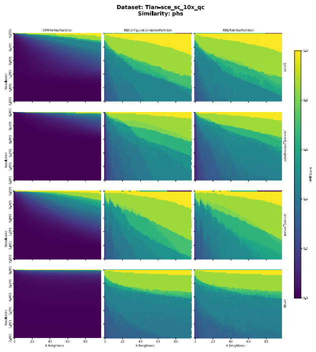


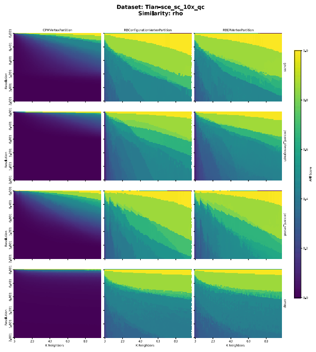


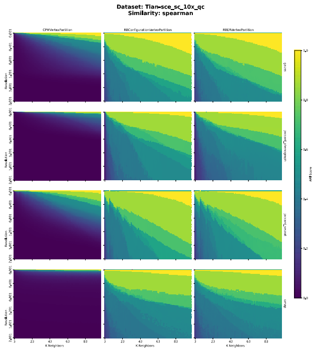


Figure 40: Performance landscapes for Tian-sce_sc_10x_qc, for: `phs`, `rho`, `spearman`. Performance landscapes illustrate the interaction between numerical hyperparameters ($k$, resolution) and categorical hyperparameters. Each 12-cell grid represents combinations of Graph Type and Quality Function for a given Similarity Measure. Color intensity encodes ARI scores, with brighter regions indicating higher clustering performance and darker regions indicating poorer results.

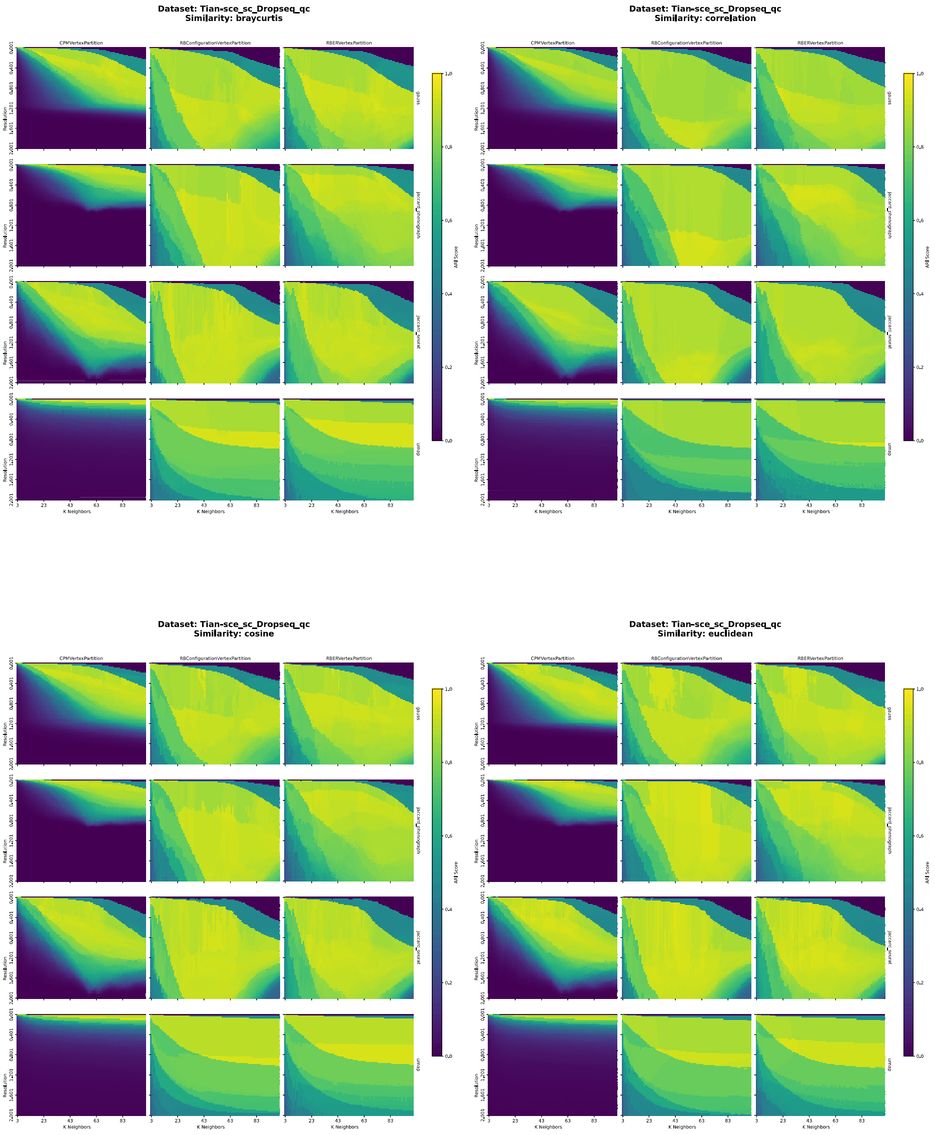


Figure 41: Performance landscapes for Tian-sce_sc_Dropseq_qc, for: `braycurtis`, `correlation`, `euclidean`, `cosine`. Performance landscapes illustrate the interaction between numerical hyperparameters ($k$, resolution) and categorical hyperparameters. Each 12-cell grid represents combinations of Graph Type and Quality Function for a given Similarity Measure. Color intensity encodes ARI scores, with brighter regions indicating higher clustering performance and darker regions indicating poorer results.

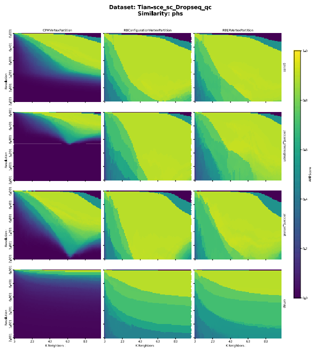


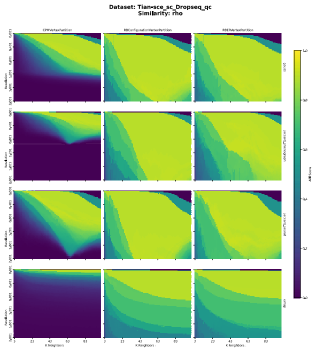


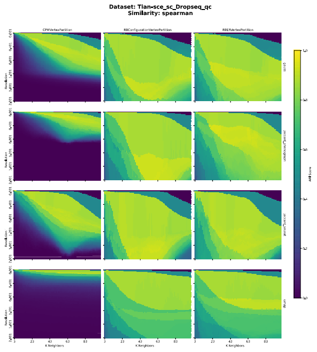


Figure 42: Performance landscapes for Tian-sce_sc_Dropseq_qc, for: `phs`, `rho`, `spearman`. Performance landscapes illustrate the interaction between numerical hyperparameters ($k$, resolution) and categorical hyperparameters. Each 12-cell grid represents combinations of Graph Type and Quality Function for a given Similarity Measure. Color intensity encodes ARI scores, with brighter regions indicating higher clustering performance and darker regions indicating poorer results.

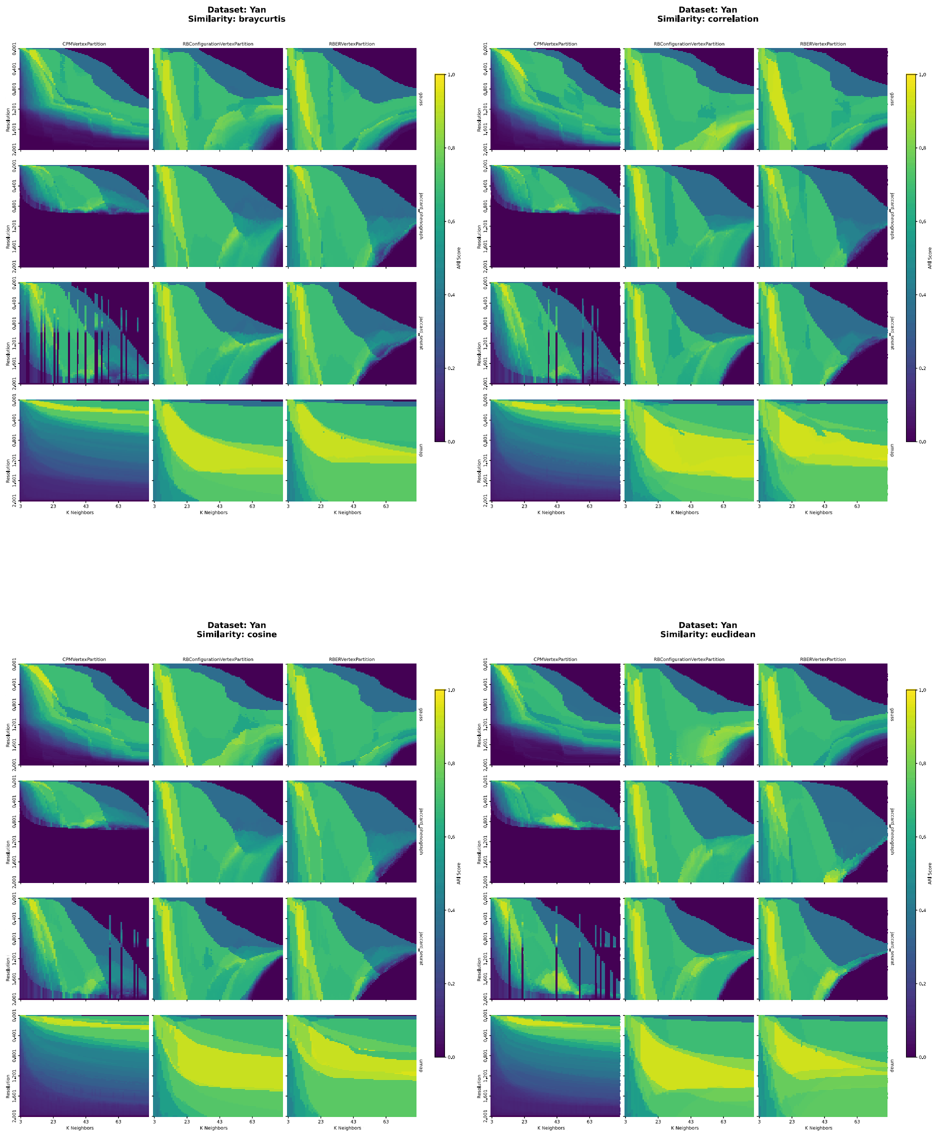


Figure 43: Performance landscapes for Yan, for: `braycurtis`, `correlation`, `euclidean`, `cosine`. Performance landscapes illustrate the interaction between numerical hyperparameters ($k$, resolution) and categorical hyperparameters. Each 12-cell grid represents combinations of Graph Type and Quality Function for a given Similarity Measure. Color intensity encodes ARI scores, with brighter regions indicating higher clustering performance and darker regions indicating poorer results.

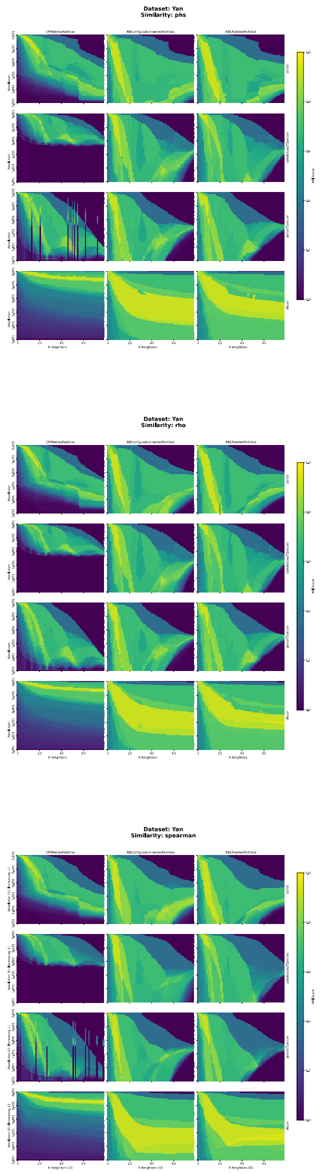


Figure 44: Performance landscapes for Yan, for: `phs`, `rho`, `spearman`. Performance landscapes illustrate the interaction between numerical hyperparameters ($k$, resolution) and categorical hyperparameters. Each 12-cell grid represents combinations of Graph Type and Quality Function for a given Similarity Measure. Color intensity encodes ARI scores, with brighter regions indicating higher clustering performance and darker regions indicating poorer results.

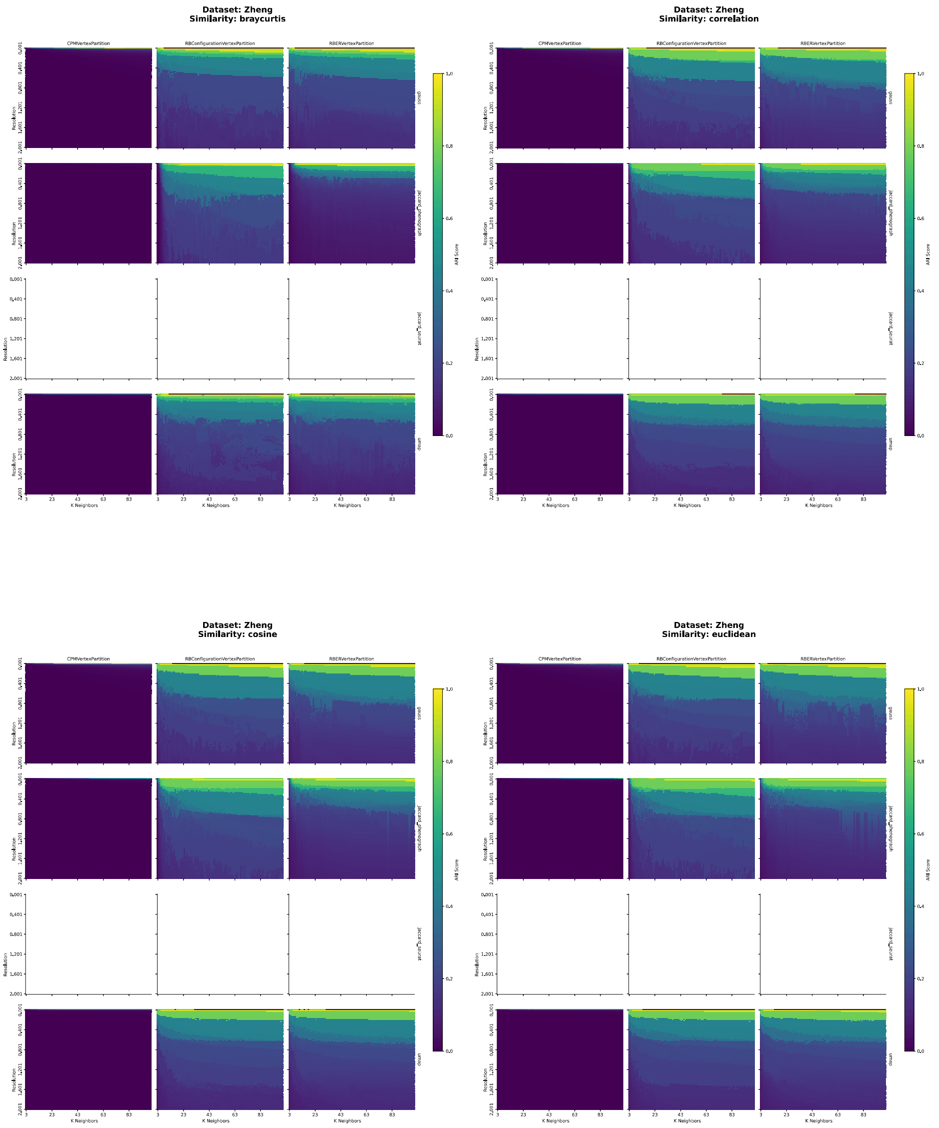


Figure 45: Performance landscapes for Zheng, for: `braycurtis`, `correlation`, `euclidean`, `cosine`. Performance landscapes illustrate the interaction between numerical hyperparameters ($k$, resolution) and categorical hyperparameters. Each 12-cell grid represents combinations of Graph Type (except for `jaccard_seurat`, which was computationally prohibitive for this dataset) and Quality Function for a given Similarity Measure. Color intensity encodes ARI scores, with brighter regions indicating higher clustering performance and darker regions indicating poorer results.

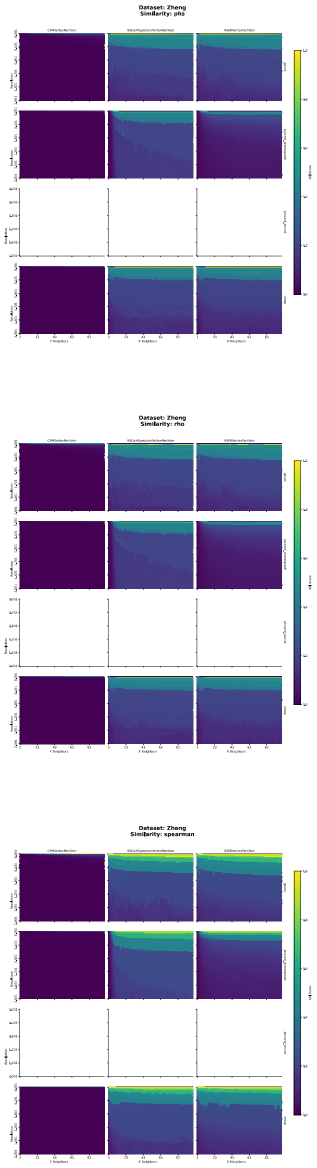


Figure 46: Performance landscapes for Zheng, for: `phs`, `rho`, `spearman`. Performance landscapes illustrate the interaction between numerical hyperparameters ($k$, resolution) and categorical hyperparameters. Each 12-cell grid represents combinations of Graph Type (except for `jaccard_seurat`, which was computationally prohibitive for this dataset) and Quality Function for a given Similarity Measure. Color intensity encodes ARI scores, with brighter regions indicating higher clustering performance and darker regions indicating poorer results.
